# Excess crossovers impede faithful meiotic chromosome segregation in *C. elegans*

**DOI:** 10.1101/2020.01.30.927640

**Authors:** Jeremy A. Hollis, Marissa L. Glover, Aleesa Schlientz, Cori K. Cahoon, Bruce Bowerman, Sarah M. Wignall, Diana E. Libuda

## Abstract

During meiosis, at least one crossover must form between each pair of homologous chromosomes to ensure their proper partitioning. However, most organisms limit the number of crossovers by a phenomenon called crossover interference; why this occurs is not well understood. Here we investigate the functional consequences of extra crossovers in *Caenorhabditis elegans*. Using a fusion chromosome that exhibits a high frequency of supernumerary crossovers, we find that essential chromosomal structures are mispatterned, subjecting chromosomes to improper spindle forces and leading to congression and segregation defects. Moreover, we uncover mechanisms that counteract these errors; anaphase I chromosome bridges were often able to resolve in a LEM-3 nuclease dependent manner, and tethers between homologs that persisted were frequently resolved during Meiosis II by a second mechanism. This study thus provides evidence that excess crossovers impact chromosome patterning and segregation, and also sheds light on how these errors are corrected.

## Introduction

Meiosis is a specialized, reductional form of cell division necessary for the production of haploid sperm or egg cells. One hallmark of meiosis is the requirement of genetic exchange through recombination. Meiotic recombination is initiated by the formation of double strand DNA breaks (DSBs), which are repaired to form crossover (CO) and noncrossover events (Gray and Cohen 2016). Germ cells in most organisms require a CO between homologous chromosomes to physically link each pair of homologs, enabling their proper segregation during the meiosis I division. Despite the formation of many programmed DSBs, most organisms are limited in the number of COs formed, and formation of a CO tends to inhibit formation of other COs nearby on the same chromosome pair, a conserved phenomenon known as CO interference (Muller 1916, Sturdevant 1913). COs that are subject to interference are considered interfering COs (or Class I COs). Additionally, some organisms have a subset of COs that are not subject to interference (called “non-interfering”, or “Class II” COs), but these non-interfering COs represent only 5-35% of all meiotic COs in *A. thaliana, M. musculus*, and *S. cerevisiae* (Gray and Cohen 2016). While CO interference is a well conserved phenomenon among most eukaryotes, the molecular consequences that result from the occurrence of multiple interfering COs are not clear. In addition, it is also unclear how these consequences may have contributed to the conservation of CO interference.

The model organism *Caenorhabditis elegans* is a particularly powerful system to study CO regulation because it exhibits remarkably strict CO control; under wild-type conditions, only one DSB per chromosome is repaired as a CO and all COs are interfering COs (Martinez-Perez and Colaiacovo 2009). Several studies have demonstrated that even in the presence of an extreme excess of DSBs (10-fold greater than wild-type levels), only a single interfering CO, marked by the pro-crossover factor COSA-1, is made per pair of homologous chromosomes (Yokoo et al. 2012, Libuda et al. 2013). Additionally, it has been shown that CO interference can operate over distances longer than the length of a normal *C. elegans* chromosome axis (Hillers and Villeneuve 2003, Libuda et al. 2013). Thus, in the case of chromosomes with an abnormally long chromosome axis, such as end-to-end fusions of chromosomes, many meioses still only have one CO per fusion chromosome pair (Hillers and Villeneuve 2003, Libuda et al. 2013).

These COs form physical connections between the homologs, known as a chiasmata. Since in *C. elegans* each chromosome pair typically has one CO that occurs off-center along the chromosome length, the chromosomes reorganize around this single chiasma to form cruciform bivalents with long and short arms (Martinez-Perez et al. 2008, Nabeshima et al. 2005). These bivalents then align on the spindle, and in anaphase I, cohesion is lost along the short arm axis, enabling segregation of homologous chromosomes (Kaitna et al. 2002, Rogers et al. 2002). Aurora B kinase (AIR-2) and other members of the conserved chromosomal passenger complex (CPC) are targeted to this short arm region in prophase to protect sister chromatid cohesion in that region until anaphase I is triggered (Martinez-Perez et al. 2008). Moreover, the CPC also directs the formation of a larger meiotic protein complex that forms upon nuclear envelope breakdown (Wignall and Villeneuve 2009, Dumont et al. 2010); at this stage, the CPC reorganizes from a linear distribution along the short arm axis to a ring encircling this region, and targets a number of other conserved proteins to form a structure known as the Ring Complex (RC). In addition to the CPC, the RC contains other conserved components, such as the kinase BUB-1 (Dumont et al. 2010) and the microtubule de-stabilizing kinesin MCAK^KLP-7^ (Connolly et al. 2015, Han et al. 2015). Furthermore, SUMO and SUMO pathway enzymes localize to the RC and are required for RC assembly and stability (Pelisch et al. 2017, Davis-Roca et al. 2018). Chromosomes that lack RCs have congression and segregation errors (Muscat et al. 2015), and depletion of RC components causes a variety of meiotic defects (Wignall and Villeneuve 2009, Dumont et al. 2010, Romano et al. 2003, Schumacher et al. 1998, Connolly et al. 2015, Pelisch et al. 2017, Laband et al. 2017, Pelisch et al. 2019), highlighting the importance of this complex.

One function of the RC is to promote chromosome congression (Wignall and Villeneuve 2009, Pelisch et al. 2017). In *C. elegans* oocyte spindles, microtubule bundles run laterally along the sides of bivalents instead of forming canonical end-on kinetochore attachments. A component of the RC, the kinesin-4 family member KLP-19, has been proposed to walk along these laterally-associated bundles towards microtubule plus ends located in the center of the spindle, thus providing chromosomes with plus-end directed forces that mediate metaphase alignment (Wignall and Villeneuve 2009). Then, at anaphase onset, the enzyme SEP-1^separase^ is targeted to the midbivalent region to cleave cohesin and allow homologous chromosomes to segregate to opposite spindle poles (Muscat et al. 2015, Siomos et al. 2001). At this stage, the RCs are removed from chromosomes and remain in the center of the spindle, where they disassemble (Dumont et al. 2010, Muscat et al. 2015, Davis-Roca et al. 2017, Mullen et al 2017, Davis-Roca et al. 2018). Oocytes extrude one set of homologs into a polar body, and then Meiosis II (MII) proceeds. The MII chromosomes assemble RCs encircling the sister chromatid interface and repeat the segregation process to form a matured haploid egg.

Although extensive research has focused on CO formation and RC function independently, it is still unclear how early meiotic processes affect chromosome structure and function in meiotic divisions. Moreover, why chromosomes in many organisms are so tightly restricted to 1-2 COs per homolog pair has not been addressed. Here we utilize a *C. elegans* strain containing an end-to-end fusion of three chromosomes to assess the effects of supernumary crossovers on meiosis. We find that the RC is mispatterned in the presence of multiple COs, leading to defects in chromosome congression, and that chromosomes with multiple chiasmata exhibit extensive chromatin bridging in anaphase. Thus, excess crossovers can severely impact chromosome patterning and segregation, highlighting the importance of limiting the number of recombination events between homologous chromosomes for the proper execution of meiosis. Further, our studies uncovered multiple mechanisms by which oocytes are able to correct these errors, demonstrating that species have evolved ways to combat the deleterious defects caused by excess crossing over.

## Results

### Fusion chromosomes generate bivalents with multiple crossovers and chiasmata

To understand the effects of multiple COs on proper chromosome segregation and gamete formation, we sought to consistently generate multiple COs along a single chromosome during meiosis. Since normal *C. elegans* chromosomes typically experience only one CO per meiosis (Hillers and Villeneuve 2003, Libuda et al. 2013), we utilized a strain containing the three-chromosome fusion *meT7 (*end-on-end fusions of chromosomes *III, X*, and *IV*; Figure 1A). While wild-type strains contain six individual chromosomes, strains containing the *meT7* fusion chromosome have four individual chromosomes total (*meT7 III; X; IV* fusion chromosome, and chromosomes *I, II*, and *V;* Figure 1B, Left). As a consequence of *meT7* being three times the length of a normal chromosome, previous studies using genetic assays found the occurrence of multiple COs along this fusion chromosome (Hillers and Villeneuve 2003).

**Figure 1.**
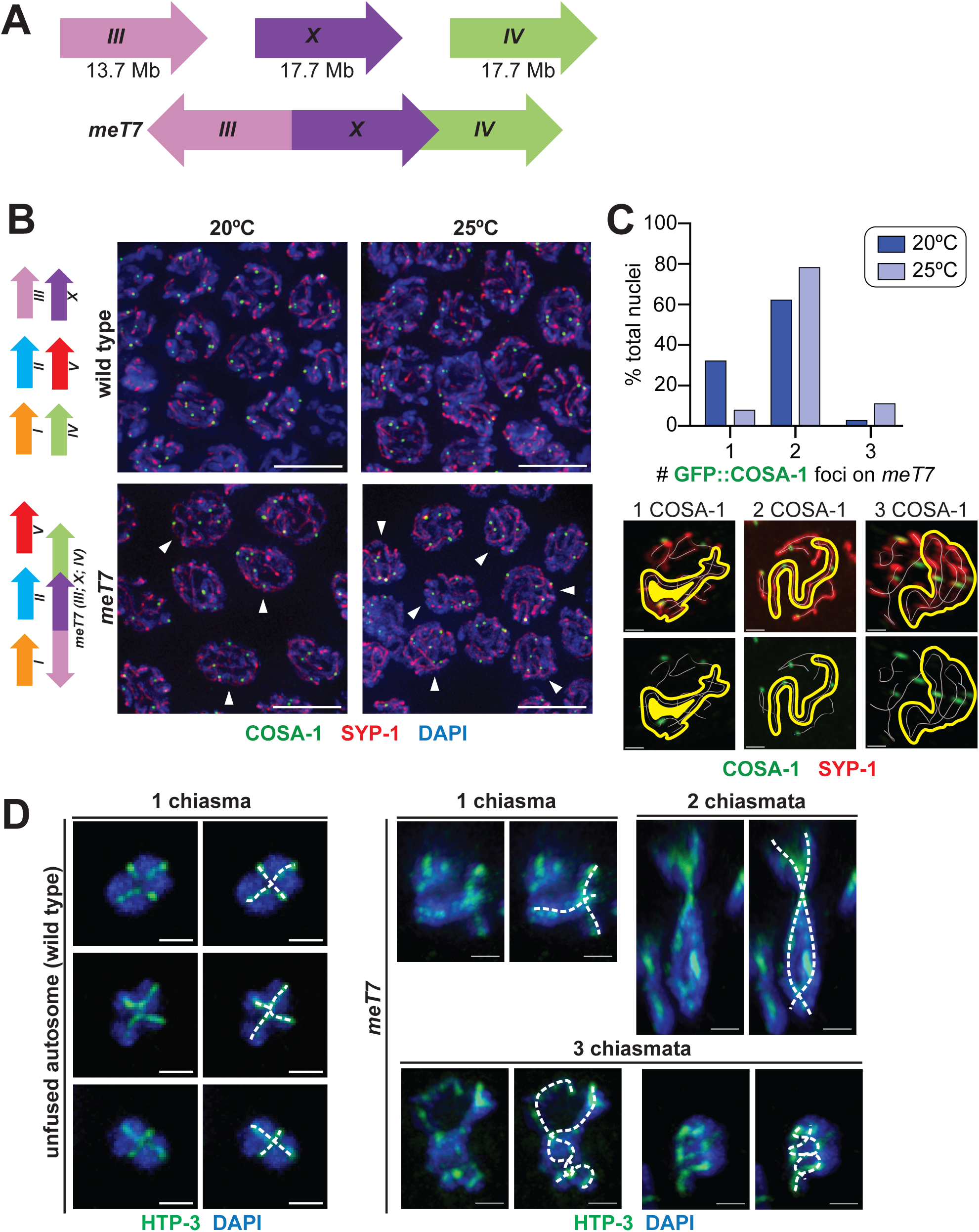
Fusion chromosomes generate bivalents with multiple crossovers and chiasmata. (A) Schematic indicating the orientation of the *meT7* fusion chromosome, which fuses the *X* chromosome and chromosomes *III* and *IV*. (B) Left, schematic depicting the chromosomes in a wild-type strain (top) and the *meT7 (X;III;IV)* fusion chromosome strain (bottom). Right, immunofluorescence images of GFP::COSA-1 in late pachytene nuclei from wild-type and *meT7* fusion chromosome strains grown at 20°C and 25°C. GFP::COSA-1 is shown in green, synaptonemal complex protein SYP-1 is shown in red, and DNA is shown in blue. Arrowheads indicate nuclei with >2 crossovers along the *meT7* fusion chromosome. Scale bars = 5 µm. (C) Top, quantitation of percentage of *meT7* nuclei with indicated number GFP::COSA-1 foci on *meT7* in late pachytene at 20°C and 25°C. Number of late pachytene nuclei scored for COSA-1 foci: 20°C, n=290; 25°C n=287. Bottom, representative immunofluorescence images of single nuclei with *meT7* chromosomes with the indicated number of COSA-1 foci. White line indicates traced chromosome axis of each chromosome in a single nucleus. Yellow outline highlights the *meT7* fusion chromosome within the nucleus. (D) Three-dimensionally rendered immunofluorescence images of individual *meT7* diakinesis bivalents. Dashed lines (white) indicate traced HTP-3 axes (green), with crossing of axes indicating chiasmata. Scale bars = 1 µm.

To determine the exact number of COs occurring along the length of *meT7* within a population of nuclei, we assessed the number of COSA-1 foci, a cytological marker of Class I interfering COs during late pachytene (Yokoo et al. 2012), along *meT7* chromosomes in single nuclei. In wild-type strains containing unfused chromosomes, the six individual chromosomes in each nucleus obtain a single CO per chromosome resulting in 6 COSA-1 foci per nucleus, with <0.4% of nuclei obtaining more than 6 foci (Yokoo et al. 2012, Libuda et al. 2013) (Figure 1B and Materials and Methods). In contrast, strains grown at 20°C with the *meT7* fusion chromosome obtain 4-6 COSA-1 foci per late pachytene nucleus (Figure 1B), with 77% of *meT7* chromosomes exhibiting ≥2 COSA-1 foci (Figure 1C). To further increase the number of COs occurring along *meT7*, we grew strains containing the *meT7* fusion chromosome at 25°C, a temperature that was previously found to increase the number of COs along the length of the two-chromosome fusion *mnT12* (Libuda et al. 2013). In contrast to 20°C, we found that *meT7* strains grown at 25°C had an increase in COSA-1 foci, with >90% of *meT7* chromosomes with 2 or more COSA-1 foci (Figures 1B and 1C). This enrichment of COSA-1 foci indicates the occurrence of multiple COs along the length of the *meT7* fusion chromosome, as previously reported (Hillers and Villeneuve 2003).

To determine whether these increased COSA-1 foci along *meT7* fusion chromosomes represent COs that become chiasmata, we assessed chiasma number and bivalent structure at diakinesis along fusion chromosome pairs. Using HTP-3 immunofluorescence to trace chromosome axes (Figure 1D), we found that 13/21 diakinesis nuclei with *meT7* bivalents at 20°C had more than one chiasma, consistent with 60-80% of the *meT7* chromosome pair having more than one COSA-1 focus at the late pachytene stage (Figure 1C). In *meT7* strains grown at 25°C, we found that 19/21 (90.5%) *meT7* fusion chromosomes had more than one chiasma, consistent with 90% of *meT7* chromosome pairs having 2 or more COSA-1 marked crossovers (Figure 1C). In accordance with previous analysis performed at 20°C (Martinez-Perez et al. 2008), we found that these multi-chiasmata *meT7* bivalents grown at 25°C exhibit unique meiotic chromosome structural reorganization (Figure 1D), which is consistent with the occurrence of multiple crossovers and altered CO patterning. COs are normally formed off-center of the *C. elegans* chromosome, resulting in the formation of long and short arms in the cruciform bivalent structure (Figure 1D). Previous studies have found that some meiotic chromosome structures reorganize around these CO sites starting at the late pachytene-diplotene transition, resulting in cytologically distinguishable long and short bivalent arms. In comparison to single chiasma *meT7* and non-fusion cruciform bivalent structures, these multi-chiasma *meT7* bivalent structures exhibit more than two short arms. Together, this corresponding increase in both COSA-1 foci and chiasmata indicates that the extra COSA-1 foci observed at pachytene in *meT7* chromosomes represent *bona fide* cytologically-differentiated meiotic CO events that can result in atypical bivalent structures.

### Bivalents with multiple crossovers have mispatterned Ring Complexes

Given these findings, we set out to investigate how extra chiasmata might affect other aspects of bivalent organization by assessing the Ring Complex (RC), a structure comprised of a set of critical meiotic proteins. AIR-2 and other CPC components localize along the short arms of each cruciform bivalent during diakinesis, and upon nuclear envelope breakdown, the CPC reorganizes into a ring encircling the short arm axis of the bivalent, and forms the RC by recruiting other components (Wignall and Villeneuve 2009, Dumont et al. 2010, Pelisch et al. 2017). To determine if this structure properly forms on bivalents with multiple short arm regions (resulting from excess COs) (Figure 1D), we compared the organization of the CPC and other RC components on wild-type and *meT7* prometaphase bivalents.

The three wild-type bivalents in the *meT7* strain formed single whole rings of AIR-2, as expected. However, while AIR-2 localized to *meT7* fusion bivalents, it was sometimes improperly shaped: 72% of *meT7* bivalents had single rings at 15°C, but other *meT7* bivalents formed either slightly (19%) or severely (10%) mispatterned structures (Figure 2A, 2B; see Materials and Methods for details on quantification). BIR-1^Survivin^, another CPC component, colocalized with AIR-2 on all ring structure types (Figure S1), suggesting that the pattern of AIR-2 localization reflects assembly of the entire CPC. Similar to the increase in COs and chiasmata at 25°C (77% *meT7* with >2 COs at 20°C versus 90% *meT7* with >2 COs at 25°C; Figure 1D), these CPC patterning defects on the *meT7* bivalent increased in number and severity at 25°C (increasing to 30% and 34%, for slight and severe mispatterning, respectively, Figure 2A), suggesting that increased CO numbers increase the likelihood of CPC mispatterning.

**Figure 2.**
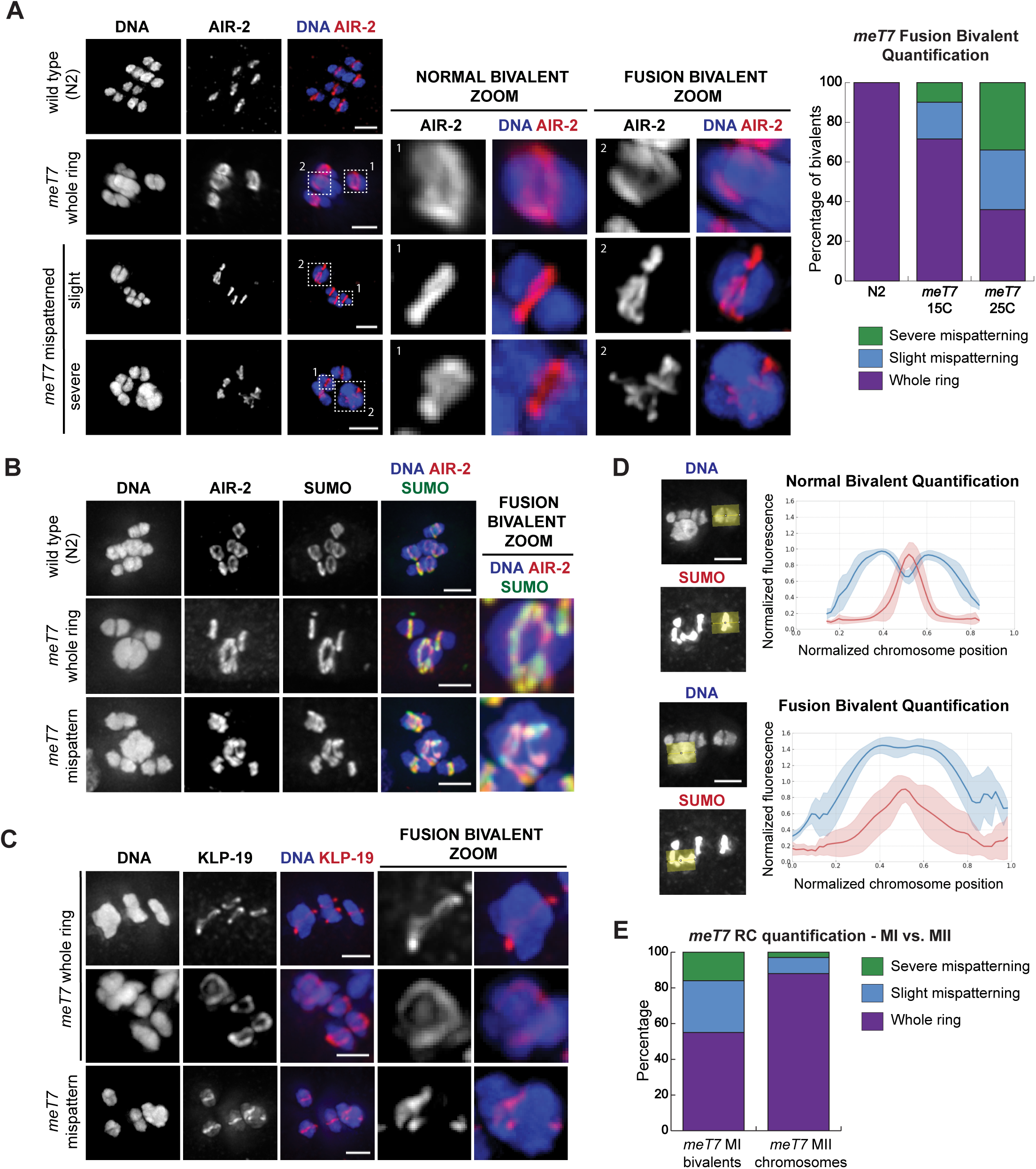
Bivalents with excess crossovers have mispatterned Ring Complexes (RCs). (A) AIR-2 localization in wild type (N2) or *meT7* oocytes. Single-chromosome zooms show that AIR-2 (red) localizes in a whole ring shape encircling normal bivalents in both strains, while its localization is either whole ring-like or mispatterned on *meT7* fusion bivalents. Note that the zoomed images throughout this figure are partial projections, chosen to highlight individual chromosomes. Quantification shows that *meT7* mispatterning increases with increased temperatures (N=75, all categories). (B) SUMO (green) co-localizes with AIR-2 (red) in all ring structure types on both wild type (N2) and *meT7* fusion bivalents. (C) KLP-19 (kinesin 4) (red), localizes to both“whole ring” and“mispatterned” *meT7* RCs. (D) Linescans across bivalents show that both the bivalent and RC have wider spread and higher variance in *meT7* fusion bivalents as compared to wild type bivalents. (N=25, all categories). (E) Quantification of ring patterning in metaphase I bivalents and metaphase II sister chromatid pairs. Metaphase I bivalents have much higher rates of mispatterning as compared to metaphase II sister chromatid pairs (N=75, all categories). All scale bars=2.5µm.

We next assessed the localization of RC components that are dependent on the CPC for targeting, and found that both SUMO (Figure 2B) and KLP-19 (Figure 2C) co-localize with AIR-2 on *meT7* bivalents. Therefore, although the CPC is mispatterned, it can still target other RC components to the bivalents. Linescans across SUMO-stained bivalents showed that, in contrast to the single tight ring peak and bilobed bivalent structure characteristic of wild-type bivalents, *meT7* bivalents have much more variation in both chromosome and RC shape (Figure 2D). Importantly, although RCs also form in Meiosis II around the interface between sister-chromatids, the RC patterning defects mainly occurred on Meiosis I *meT7* bivalents (45% of all MI *meT7* bivalents), while only 12% of MII chromosomes formed abnormal rings (p<0.0001, chi-square test; Figure 2E). This result is consistent with the idea that the RC defects are primarily caused by excess COs between homologous chromosomes, rather than general problems with RC formation in the *meT7* strain.

Although the RCs were mispatterned on *meT7* bivalents, kinetochore proteins appeared to load on the bivalents. Since *C. elegans* chromosomes are holocentric, kinetochore proteins coat the bivalents in meiosis (Howe et al. 2001, Monen et al. 2005). We found that two kinetochore components, BUB-1 and SEP-1^Separase^, coated the entire fused bivalent, demonstrating that the major patterning defect on fusion chromosomes was with the RCs, not the kinetochore (Figure S2).

### Fusion chromosome bivalents can form multiple Ring Complexes that all remain functional

Next, we wanted to characterize the mispatterned *meT7* RC structures. In some of our images there appeared to be multiple distinct RCs forming on a single *meT7* bivalent (e.g. Figure 2A, row 3; Figure 2B, row 3), which could reflect the ability of each chiasmata to organize its own RC. However, since many of the mispatterned RCs appeared to be complex structures, it was difficult in many cases to determine if a particular RC was comprised of multiple rings close together, or instead represented a single intertwined structure. Therefore, to distinguish between these possibilities, we performed an “RC stretching assay.” This assay exploits our previous finding that under extended metaphase arrest, RCs begin to stretch away from the chromosomes towards microtubule plus ends, reflecting the fact that they contain a plus-end-directed activity that usually provides chromosomes with plus-end-directed forces (Muscat et al. 2015). For the purposes of the current study, we reasoned that this behavior might spatially separate distinct rings from one another, enabling us to distinguish and quantify the total number of rings formed on each *meT7* fusion bivalent. Additionally, we performed this “RC stretching assay” on monopolar spindles, in which the microtubule minus ends are organized at a central pole and the plus ends radiate outward forming an aster. This additional feature enabled us to determine if the *meT7* ring structures were functional (*i.e.* whether they were able to stretch towards the outside of the aster, thus exerting plus-end forces). Consistent with previous work (Muscat et al. 2015), we found that in this RC stretching assay, rings on normal bivalents typically stretch off as one entity in a single direction towards microtubule plus ends (31/38), with 7/38 of the observed bivalents having two stretches (Figure 3, rows 1 and 2). In contrast, on many *meT7* bivalents, it was clear that there was more than one RC, often with two (27/38) or rarely three (2/38) separate entities stretching off of the bivalent (Figure 3, rows 5 and 6). Notably, the frequency of more than one stretching RC on the *meT7* fusion chromosome (76% at 15°C) is close to the frequency of more than one CO along *meT7* (77% at 20°C; Figure 1C). This result suggests that multiple distinct RC structures can form at different chiasmata on the same bivalent.

**Figure 3.**
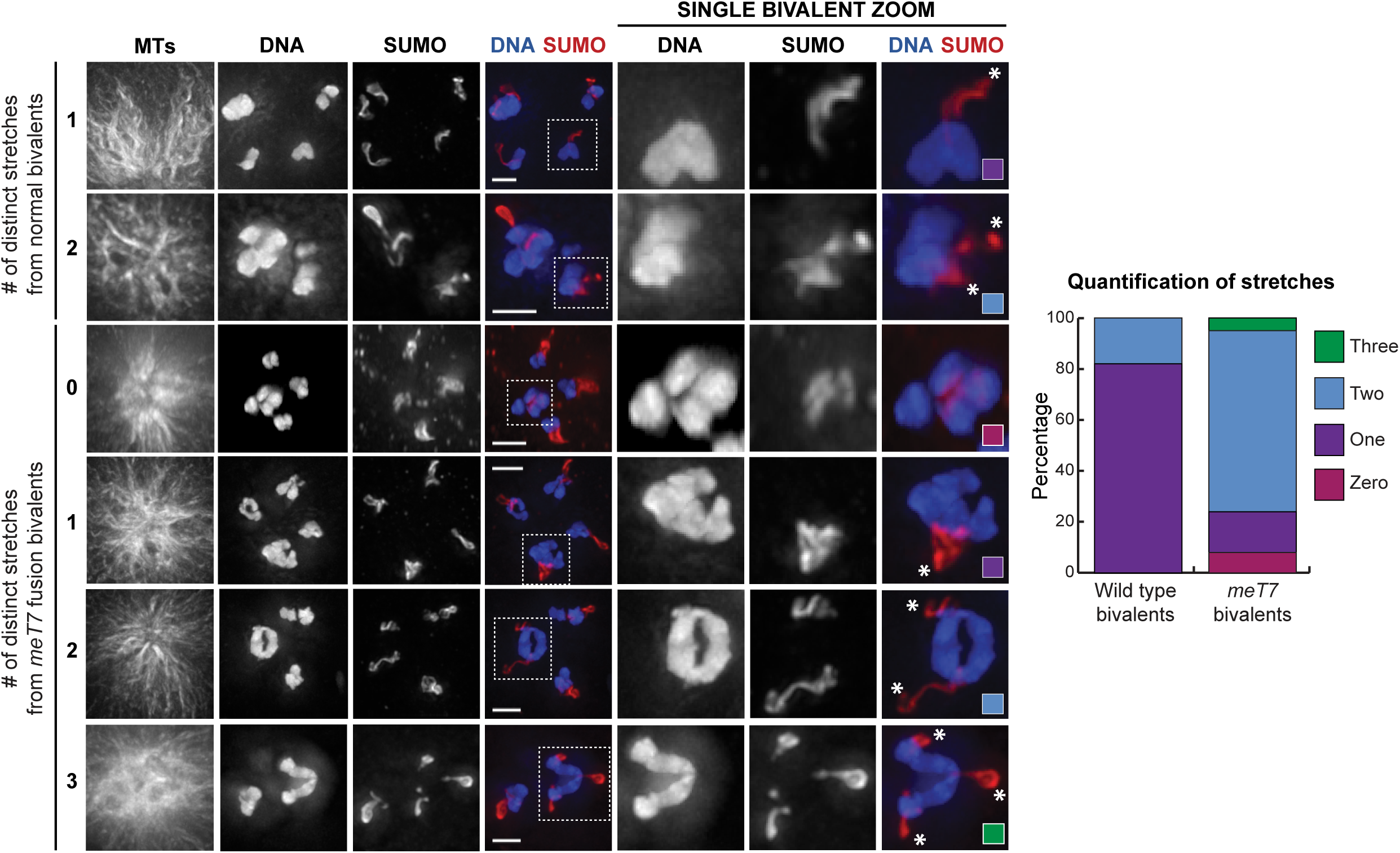
Bivalents with excess COs can form multiple functional RCs. Examples and quantification of stretching RCs (visualized by SUMO staining, red) from bivalents (blue) (N=38, all conditions). In this assay, monopolar spindles were generated by depleting the force-generating motor KLP-18 (Wignall and Villeneuve 2009, Wolff *et al.* 2016). Zooms are partial projections, chosen to highlight individual chromosomes. RCs on monopolar spindles under prolonged metaphase arrest tend to stretch in a plus end-directed manner, towards the outside of the microtubule aster. RCs on normal bivalents tend to stretch as mostly one unit in a single direction, while RCs on *meT7* bivalents can have multiple RCs that stretch in independent directions. Asterisks denote stretching RCs. Scale bars=2.5µm.

Interestingly, when multiple RCs are present, they all stretch towards microtubule plus ends but end up oriented along different microtubule bundles, suggesting that the RCs are all functional and acting independently of one another; in the context of a bipolar spindle where microtubule bundles are not all in the same orientation, this could result in a single chromosome being pulled in opposite directions. Furthermore, unlike on normal bivalents, 3/38 rings on *meT7* bivalents were not stretching in any direction despite normal bivalents in the same spindle having stretching rings. Together, these results suggest that the mispatterned RCs on *meT7* may not be able to provide chromosomes with normal plus-end-directed forces.

### Fusion chromosome bivalents show defects in metaphase alignment and bivalent organization

To investigate the possibility that *meT7* bivalents are experiencing abnormal forces, we asked whether mispatterning of the RCs in the *meT7* bivalents had functional consequences on the alignment of those bivalents on bipolar spindles. We therefore evaluated spindles where the three normal bivalents had aligned at the metaphase plate and scored the position of the *meT7* fusion bivalent. We found that while *meT7* bivalents with single whole rings aligned with the other bivalents in 92% (46/50) of these spindles, only 62% (31/50) with mispatterned rings successfully aligned (Figure 4A), suggesting that defects in RC organization affect the fidelity of chromosome congression.

**Figure 4.**
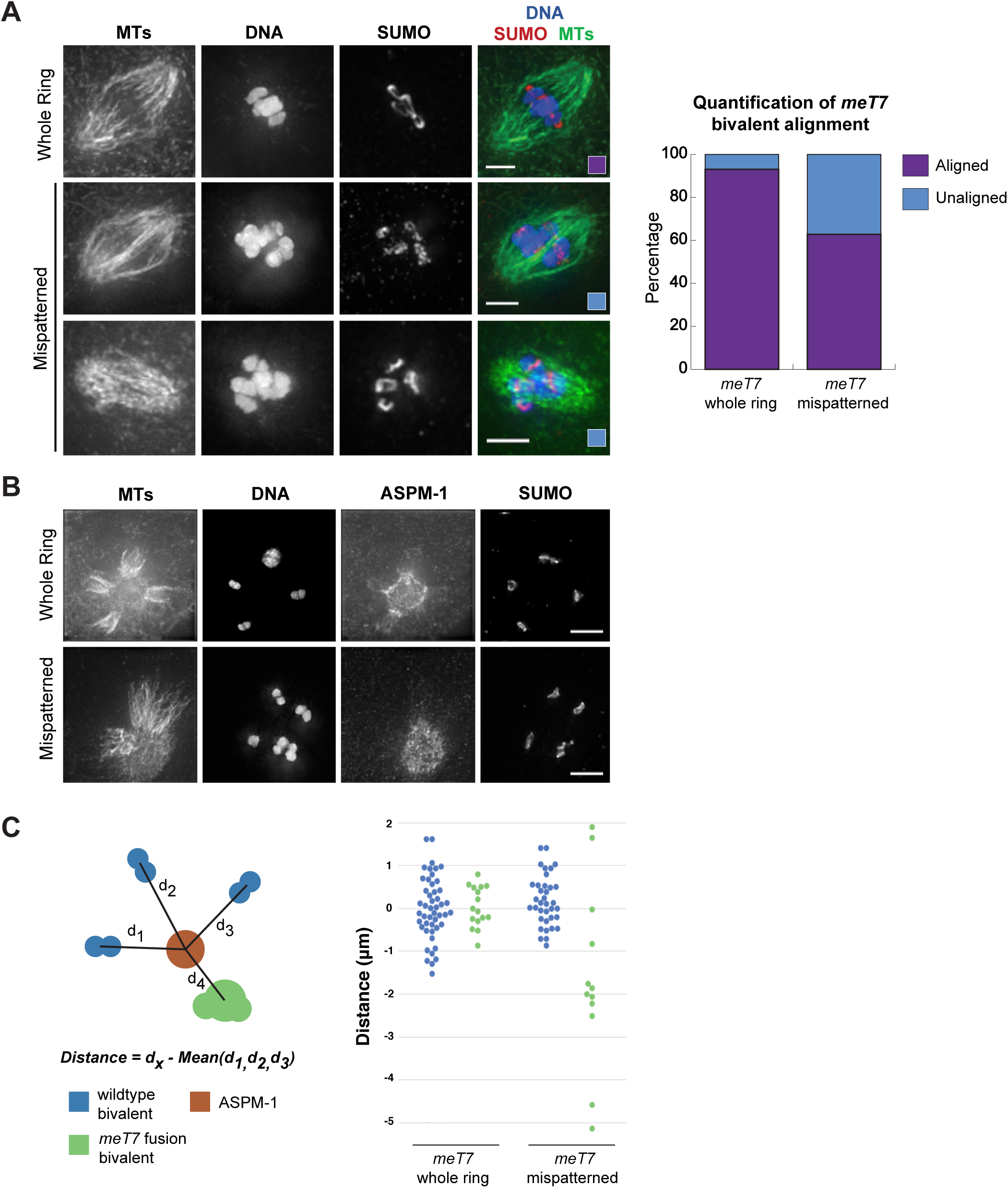
Bivalents with mispatterned RCs align improperly on the oocyte spindle and are not subjected to proper plus-end directed forces. (A) Alignment of the fusion chromosome was assessed on Metaphase I spindles; images show DNA (blue), microtubules (green) and SUMO to mark the RCs (red), and the quantification is to the right of the images. 92% of *meT7* bivalents with a whole ring aligned with the three normal bivalents, compared to 62% of *meT7* bivalents with mispatterned rings (N=50, all conditions). Scale bars=2.5µm. (B,C) Examples and quantification of plus-end directed movement of fused chromosome bivalents. Monopolar spindles were generated by depleting KLP-18, and the center of the aster was determined by staining with spindle pole marker ASPM-1 and determining the center of the ASPM-1 signal. The distance between each of the three normal bivalents and the center of the monopolar spindle was measured using Imaris, and the distance the fused chromosome bivalent traveled was compared to the average of the three normal bivalents’ distance. *meT7* bivalents with whole rings (N=16) tended to move as far as normal bivalents on the monopole, while bivalents with mispatterned rings (N=12) showed much more variation in their distance from the monopole. Scale bars=5µm.

Building on this finding, we assessed the position of chromosomes on monopolar spindles as a read-out of plus-end directed forces (Figure 4B); in this context, bivalents that are able to generate normal plus-end forces migrate away from the center of the aster (Wignall and Villeneuve 2009). Unlike the RC stretching assay with monopolar spindles (Figure 3), this experiment does not involve an extended metaphase arrest so the RCs retain their original morphology (Figure 4). This analysis revealed that *meT7* bivalents with mispatterned rings do not tend to migrate as far away from the pole as the three wild-type bivalents (Figure 4C). Moreover, in two extreme cases (out of 12), the fusion bivalents remained stuck at the center of the monopole (Figure 4B, bottom row). Importantly, this defect appeared to be caused by the chromosome patterning defects and not by the large size of the fusion bivalent, since *meT7* bivalents with single rings exhibited normal chromosome movement (Figure 4B, 4C). Together, these findings suggest that the mispatterned RCs on *meT7* bivalents are not able to provide these bivalents with normal plus-end forces, potentially impacting their ability to achieve proper metaphase alignment. Since *meT7* bivalents often have multiple RCs that can function independent of each other (Figure 3), we postulate that these RCs could provide bivalents with forces in opposing directions thereby reducing the efficiency of chromosome movement.

### Fusion chromosome bivalents show defects in chromosome segregation that carry over into Meiosis II

Given the defects in the structure and alignment of *meT7* bivalents, we next wanted to determine if having excess COs impacts chromosome segregation during meiosis I and II. First, we imaged anaphase I to assess the ability of homologous chromosomes to segregate. Notably, we found that in early anaphase I, the *meT7* fusion bivalent was often stuck in the middle of the spindle, lagging behind the three segregating wild-type bivalents. However, the frequency of this delayed segregation did not appear to be affected by temperature, as *meT7* bivalents showed comparable segregation delays in early anaphase I spindles at both 15°C (44/75, 58.7%) and 25°C (47/75, 62.7%) (Figure 5A). In contrast, in late anaphase I *meT7* bivalents had chromatin bridges that increased at higher temperatures where *meT7* experiences elevated COs (17/75, 22.7% at 15°C vs. 34/75, 45.3% at 25°C; P = 0.003, chi-square test), while wild-type bivalents segregated normally at all temperatures (Figure 5A). These results suggest that the early anaphase I defects are not due to increases in CO numbers, and may instead be due to the large size of the fusion chromosome. Conversely, extra COs between homologous chromosomes likely cause chromatin bridges in late anaphase I. Consistent with this hypothesis, we rarely observed chromatin bridging in anaphase II when sister chromatids rather than homologous chromosomes were segregating (2/65, 3.1%; Figure 5C).

**Figure 5.**
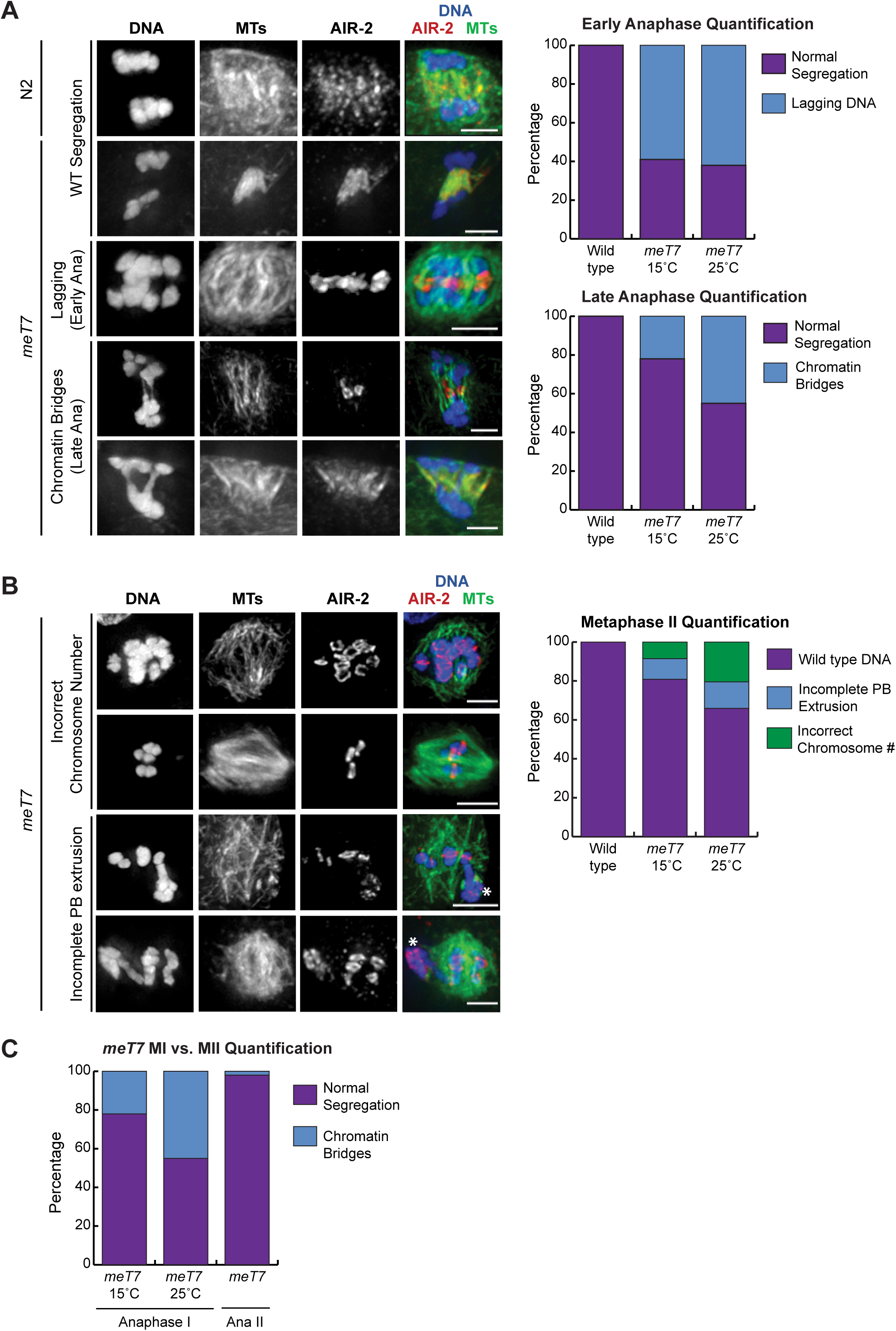
Fusion chromosome segregation is frequently aberrant. (A) Anaphase I chromosome segregation in wild-type (N2) and *meT7* oocytes; shown are DNA (blue), microtubules (green), and AIR-2 (red), with quantification to the right of the images. All chromosomes segregated without errors in wild type (N2) oocytes (rows 1 and 2), but *meT7* bivalents tended to show a delay in segregation in early anaphase compared to normal bivalents at both 15°C and 25°C (row 3, “early anaphase quantification” graph). In mid-to-late anaphase, 23% of *meT7* bivalents showed extended chromatin bridging at 15°C, which increased to 45% at 25°C (rows 4 and 5, “late anaphase quantification” graph (N=75, all conditions). (B) Metaphase II spindles in *meT7* oocytes. While all N2 metaphase II spindles were euploid, at 15°C 8% of *meT7* spindles were aneuploid (rows 1 and 2) had 12% had DNA tethered to the first polar body (indicated with asterisks, rows 3 and 4). These numbers increased to 15% and 20%, respectively, at 25°C (N=60, all conditions). (C) Quantification of *meT7* chromatin bridging in anaphase I vs. anaphase II. 22% of mid-to-late *meT7* anaphase I spindles at 15°C (N=71) and 45% at 25°C (N=58) showed anaphase chromatin bridging, while only 3% of all *meT7* anaphase II spindles (both temperatures combined) showed anaphase chromatin bridging (N=65). All scale bars=2.5µm.

To determine if the anaphase I defects had lasting consequences, we examined oocytes that progressed to Meiosis II. Notably, we observed a range of severe defects at 15°C, including oocytes that had a prominent DNA bridge connecting the *meT7* chromosome to the first polar body (5/60), as well as oocytes that had completely retained or lost the entire *meT7* bivalent, resulting in aneuploidy (7/60, Figure 5B). Similar to the chromosome bridging observed in MI, these defects increased at higher temperatures (at 25°C, 9/60 were tethered to PBI and 12/60 were aneuploid), suggesting that they were the result of excess COs between *meT7* chromosomes.

To understand the temporal dynamics of *meT7* segregation, we performed live imaging using a *meT7* strain expressing GFP-tubulin and mCherry-histone to mark microtubules and chromosomes, respectively. Consistent with our fixed imaging, we found that a majority of anaphase I spindles displayed extended chromatin bridges (8/14, 57%) (Figure 6). Five of these bridges were unresolved, leaving the first polar body tethered to the developing meiosis II spindle (Movie S1). Further, in one example, *meT7* was not able to segregate in anaphase I, and the fully retained bivalent segregated with a chromatin bridge in anaphase II (Movie S2). Altogether, this live imaging data supports the conclusion that oocytes face severe chromosome segregation defects in the presence of supernumerary crossovers.

**Figure 6.**
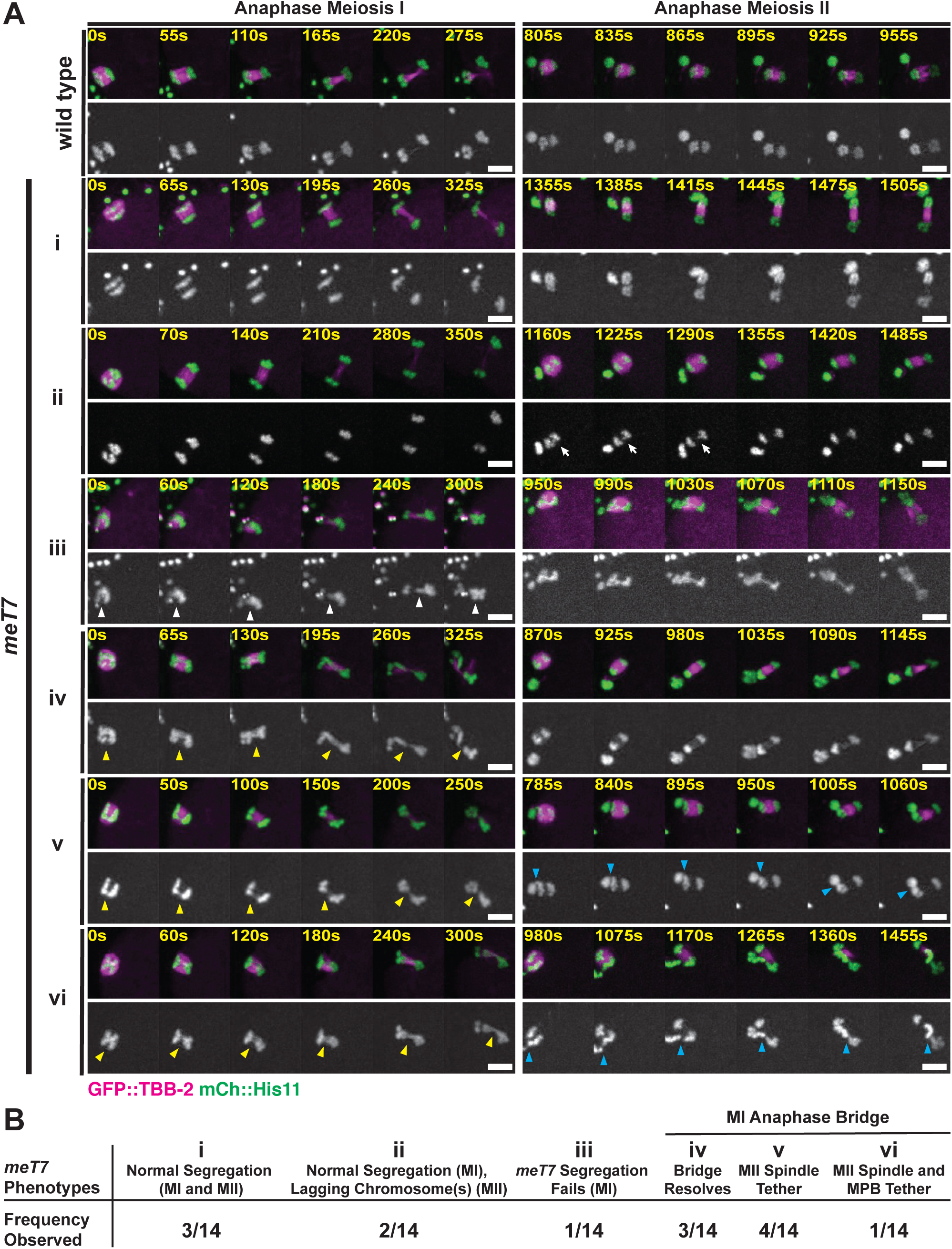
*meT7* oocytes display several classes of chromosome segregation defects. (A) Time-lapse spinning disk confocal montages during anaphase meiosis I (MI) and anaphase meiosis II (MII) in live wild-type and *meT7* oocytes expressing GFP::TBB-2 and mCherry:HIS-11 to mark microtubules and histones, respectively. Examples of *meT7* chromosome segregation phenotypes shown in i-vi. White arrowheads point to *meT7* fused chromosomes as they fail to segregate in anaphase I (iii), yellow arrowheads indicate anaphase I *meT7* chromatin bridges between segregating chromosomes (iv, v, vi), and blue arrowheads point to chromatin tethers between the MI polar body (MPB) and the anaphase II spindle (v, vi). White arrows indicate lagging chromosome(s) in anaphase II (ii). Time zero is the time at which chromosome segregation appears to begin. Scale bars, 5µm. (B) *meT7* chromosome segregation phenotype descriptions and observation frequency. Of the 8 *meT7* oocytes that had chromatin bridges at anaphase I, 3/8 bridges were able to resolve prior to the end of anaphase I. For the remaining 5 *meT7* oocytes that were not able to resolve their anaphase I bridges, 4/8 extruded the tether (with the associated chromatid) into the MPB at anaphase II and 1/8 exhibited persistent tethers through anaphase II.

### Persistent chromatin bridges can be recognized and resolved by the oocyte

Despite the fact that we observed a variety of severe meiotic chromosome segregation defects (Figures 5 and 6), the embryonic lethality of the *meT7* strain is surprisingly low (Table 1). The *meT7* strain exhibits 27.6% embryonic lethality at 25°C, indicating that most oocytes are able to generate viable progeny even in the presence of multiple COs (at 25°C, >90% of *meT7* chromosomes have ≥2 COSA-1 foci; Figure 1C) and chromosome segregation defects (at 25°C, ∼50% of *meT7* anaphase I nuclei have chromatin bridges; Figures 5 and 6). Similarly, Meiosis II oocytes have fewer defects than would be expected given the frequency of errors in Meiosis I, and in our live imaging we noticed instances where anaphase I DNA bridges were resolved as the oocyte progressed to Meiosis II (Figure 6, Movie S3). Thus, oocytes appear to have mechanisms to correct and resolve chromosome segregation defects. Supporting this idea, *meT7* oocytes appear to detect errors. We recently demonstrated that *C. elegans* oocytes recognize meiotic errors and respond by delaying AIR-2 relocalization to the spindle in mid anaphase (Davis-Roca et al. 2017). Thus, we assessed AIR-2 localization and found that *meT7* oocytes delay AIR-2 relocalization at higher rates than wildtype oocytes at both 15°C (3/31, 9.7%, in wild type vs. 7/31, 22.6% in *meT7*) and at 25°C (14/42, 33.3% in wild type vs. 12/29, 41.3% in *meT7*), further suggesting that these oocytes are responding to and attempting to resolve error (Figure S3).

**Table 1.**
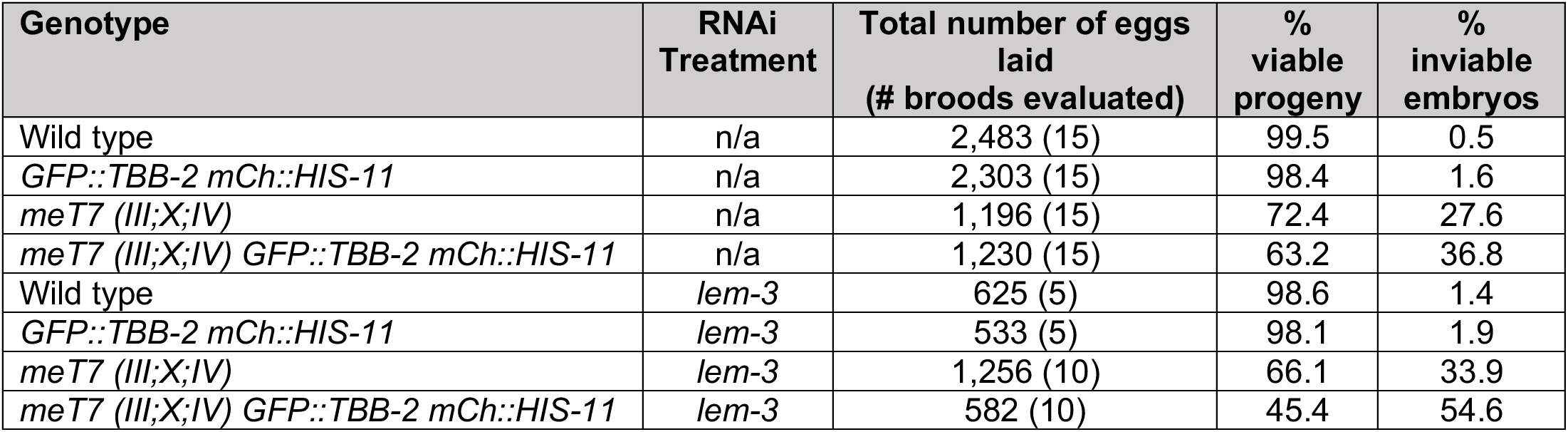
Progeny viability following lem-3 RNAi treatment.

One protein that we hypothesized might contribute to error correction is LEM-3, a late-acting nuclease proposed to resolve chromatin connections in mitotic and meiotic anaphase (Hong and Sonneville et al. 2018, Hong and Velkova et al. 2018). Therefore, we depleted *lem-3* with RNAi in the *meT7* strain and assessed chromosome segregation. In support of our hypothesis, we found that *lem-3*-depleted *meT7* oocytes had significantly higher proportions of anaphase I bridges at both 15°C (15/52, 28.8%, in control vs. 26/52, 50.0%, in *lem-3 RNAi;* P = 0.024, chi-square test) and 25°C (25/49, 51.0%, in control vs. 27/37, 72.97%, in *lem-3 RNAi*; P = 0.039, chi-square test; Figure 7A). Moreover, these defects persisted into Meiosis II in higher proportions compared to oocytes without *lem-3* depletion, and MII spindles had increased instances of both aneuploidy (at 15°C, 4/48 in control vs. 4/31 in *lem-3 RNAi, P =* 0.006, chi-square test; at 25°C, 6/41 in control vs. 6/28 in *lem-3 RNAi,* P=0.015, chi-square test) and tethered polar bodies (at 15°C, 5/48 in control vs. 12/31 in *lem-3 RNAi;* at 25°C, 9/41 in control vs. 14/28 in *lem-3 RNAi*, Figure 7A). Live imaging of *lem-3* depleted *meT7* oocytes corroborated these findings, showing an increase in chromosome segregation defects, including two cases in which chromatin bridges appeared to fragment (2/13, 15%), and one failure to extrude a Meiosis I polar body (1/13, 7%) (Figure 7B; Table S1). In five cases, the *meT7* chromosome was unable to segregate in anaphase I and subsequently extruded into the first polar body (5/13, 38%) (Figure 7B). Together, these results suggest that LEM-3 plays a role in correcting *meT7* chromosome segregation defects during oocyte meiosis.

**Figure 7.**
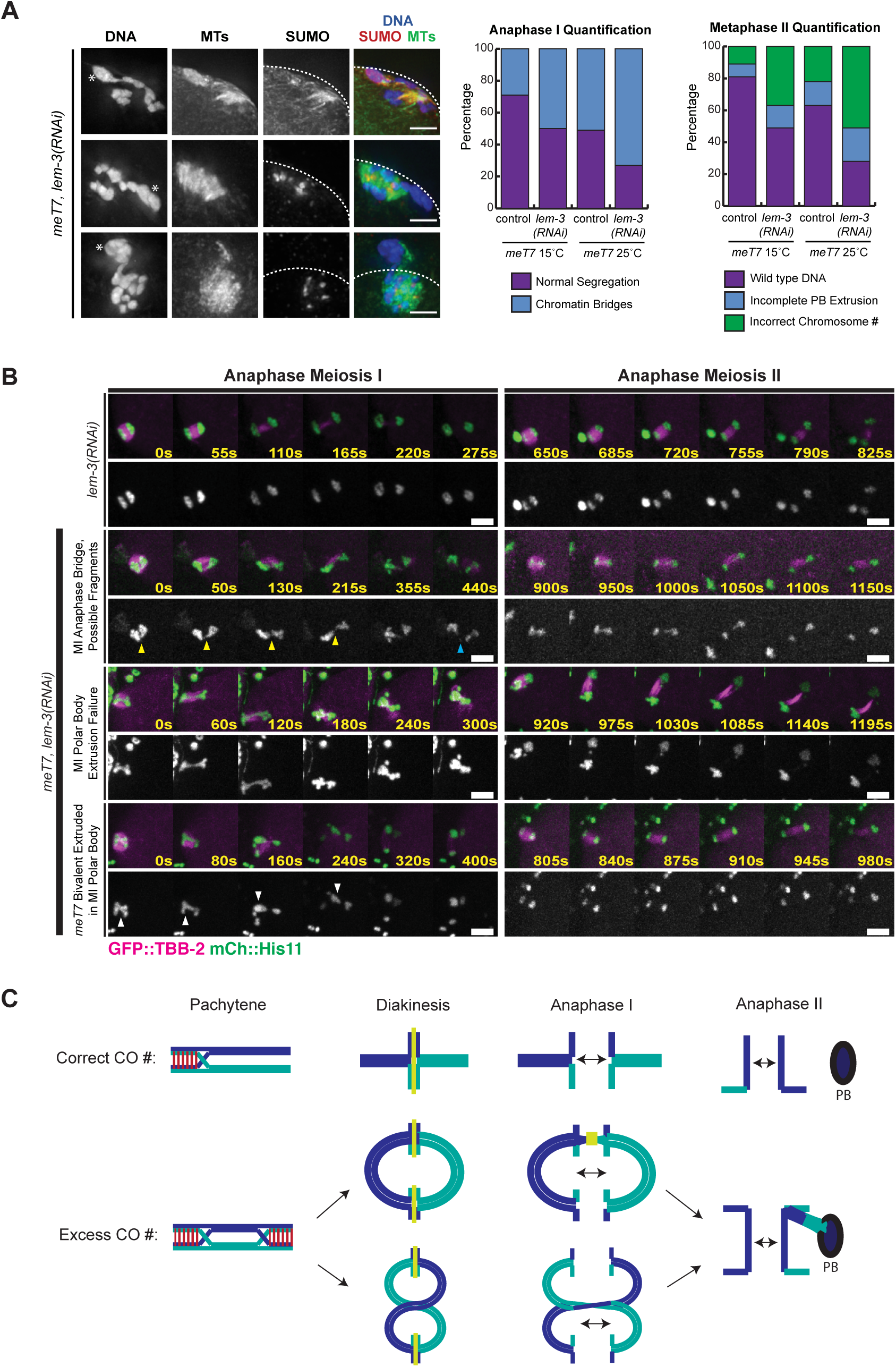
Oocytes can correct errors from excess COs in anaphase. (A) Effects of *lem-3* RNAi on *meT7* anaphase I and metaphase II spindles; shown are DNA (blue), microtubules (green) and SUMO (red). Frequency of anaphase I bridges on *meT7* increased following *lem-3*(RNAi) at both 15°C (29% for control compared to 50% for *lem-3*(RNAi); P = 0.024, chi-square test; N=52 for each condition) and 25°C (51% compared to 73%; P = 0.039, chi-square test; N=49 for control and N=37 for *lem-3*(RNAi)). Frequency of polar body extrusion delays, scored in *meT7* metaphase II spindles, increased at both 15°C (11% to 39%; P = 0.006, chi-square test; N=48 for control and N=31 for lem-3(RNAi)) and 25°C (22% to 50%; P=0.015, chi-square test; N=41 for control and N=28 for *lem-3*(RNAi)), as did aneuploidy in meiosis II at 15°C (8% to 13%) and 25°C (15% to 21%). In 23/26 of *lem-3*(RNAi) oocytes in which the polar body tether persisted into anaphase II, the tethered sister chromatid appeared to be segregating to the cell cortex (three examples shown, polar body denoted with asterisks). Scale bars=2.5µm. (B) Time-lapse montages of anaphase I and II in *lem-3* depleted wild type and *meT7* oocytes expressing GFP::TBB-2 (microtubules) and mCherry::HIS-11 (histones). Yellow arrowheads indicate an anaphase I chromatin bridge. Blue arrowhead denotes a possible chromosome fragment after bridge breaking. White arrowheads point to a failure to segregate *meT7* in Meiosis I and extrusion into the first polar body. Time zero is the time at which chromosome segregation appears to begin. Scale bars, 5µm. (C) Model for effects of supernumerary crossovers in *C. elegans* meiosis.

Furthermore, our analysis suggested another potential mechanism that could contribute to error correction in cases of chromatin bridges caused by excess COs. Specifically, we noticed that when oocytes had polar body tethers that persisted into anaphase II, the segregating sister chromatid connected to the polar body tether was often segregated to the cortical side (23/26 cases; Figure 5C), meaning that the tether was expelled with the second polar body, leaving the oocyte euploid. This fixed imaging result is supported by our live imaging, where we found that in 4/5 movies containing persisting tethers in anaphase II (Figure 6A v-vi, 6B v-vi), the sister chromatid associated with the chromosome tether was extruded with the second polar body during Meiosis II (Figure 6A v, 6B v). Currently, it is unclear whether the oocyte actively recognizes the tether or whether the DNA tether biases the direction of spindle rotation towards the cortex, thereby facilitating its own elimination. In either case, this mechanism could serve to eliminate improperly segregated chromosomes from the resulting gamete.

## Discussion

In summary, our work provides molecular insights into how crossover interference promotes proper chromosome congression and segregation in the oocyte. We found that increased COs cause defects in bivalent organization that impact the ability of chromosomes to achieve metaphase alignment. An increased number of COs leads to the designation of multiple short arms and subsequent formation of extra RCs (Figure 7C). Interestingly, when multiple RCs form along a single chromosome, they all appear to be functional. This result is in contrast to dicentric chromosomes in mice, which have multiple centromeres yet have evolved a method to inactivate all but one of them, making the chromosomes functionally monocentric (Stimpson et al. 2012). The inability to inactivate extra RCs in *C. elegans* meiosis appears to have consequences, as there are chromosome congression errors that likely arise from differential forces exerted on bivalents. This result highlights the importance of proper RC patterning, and demonstrates that the structure of this protein complex affects its function. Moreover, our studies revealed that increasing CO number has consequences for homolog segregation, as bivalents with excess crossovers have frequent chromatin bridging during the first meiotic division. These frequently observed chromosome bridges may be due in part to the formation of extra RCs along a single chromosome, which may cause chromosome alignment issues along the metaphase I plate (Figure 7C).

### Crossover limitation promote accurates chromosome segregation

Strong CO interference, where there is only ∼1-3 COs per chromosome pair per meiosis, is observed in several model systems, including fruit flies, worms, and mice (Gray and Cohen 2016). Even in human oocytes, CO interference is strong, with 1-2 COs per homolog (Wang et al. 2017). Although CO interference was initially observed over a century ago in *Drosophila* and subsequently found to be widely conserved among many model systems, the consequences of extra interfering COs along a pair of homologous chromosomes has not been directly determined. Our study is the first, to our knowledge, in any organism to assess these consequences. Our findings in *C. elegans* suggest that CO interference helps promote proper meiotic chromosome alignment by limiting homolog pairs to one crossover per bivalent under normal circumstances (thereby preventing the difficulties that ensue when bivalents assemble multiple RCs). Moreover, CO limitation may promote accurate chromosome segregation by creating a chromosome structure that facilitates the loss of cohesion in the correct domain, thereby preventing chromosome bridging. Thus, the ability to limit crossovers is important for the faithful execution of meiosis, which may represent a driving force that has contributed to the conservation of CO interference in many organisms. In the future, it will be interesting to investigate how supernumary COs affect meiotic chromosome segregation in other organisms.

In comparison to humans and other model organisms, *C. elegans* exhibits extremely strong CO interference known as complete CO interference, where one (and only one) CO is formed per chromosome pair (Martinez-Perez and Colaiacovo 2009). One potential reason why CO interference may be so robust in *C. elegans* is that they have holocentric chromosomes, which assemble centromere and kinetochore proteins along their entire length. In comparison to monocentric organisms, most holocentric organisms experience complete CO interference, suggesting that there is a strong selection bias for this highly stringent form of CO interference in these organisms (Nokkala et al. 2004, Wendel et al. 2012).

In contrast to monocentric chromosomes, holocentric chromosomes are faced with the unique challenge that microtubules could theoretically attach to any part of the bivalent surface, rather than associating in a manner that promotes the biorientation of homologous chromosomes (Melters et al. 2012). One way that *C. elegans* counteract this problem is the asymmetric positioning of a single crossover along the length of a chromosome; this generates a single point of organization along the length of each chromosome, which patterns the chromosome in a manner that facilitates chromosome biorientation, alignment, release of cohesion, and segregation (Melters et al. 2012). Perhaps the acquisition of holocentric chromosomes requires extreme crossover interference in addition to asymmetric placement of a crossover along the chromosome length. Future experiments or analysis of the evolution of holocentric organisms may parse out the correlation between holocentricity and strong crossover interference.

### Licensing of Ring Complex formation by crossovers

Our results further establish how the coordination and interplay between recombination and chromosome segregation are critical for the maintenance of genomic integrity through generations. We observed that the occurrence of multiple crossovers leads to the formation of multiple RCs. Moreover, we found a strong correlation between CO number and eventual RC number along a single chromosome. This result leads to the compelling hypothesis that a CO site may be a licensing event for RC formation. Through either its specific DNA conformation or the recombination machinery associated with it, a CO may designate a location along the chromosome for RC components to assemble. Previous experiments have already found that Aurora B kinase localizes to sites of COs (Martinez-Perez et al. 2008). Further, the HIM-6/BLM helicase has been found to localize to interfering CO sites and helps maintain chiasmata until metaphase I (Hong et al. 2016, Schvartzstein et al. 2014). Future experiments exploring whether there is direct interaction between recombination machinery and RC components may establish this connection between CO sites and RC formation.

### Mechanisms exist to correct errors caused by supernumary crossovers

Our studies also uncovered two mechanisms that counteract defects caused by excess COs. We found that depletion of the nuclease LEM-3 increases the frequency and persistence of anaphase bridging, consistent with its previously proposed role in processing erroneous recombination intermediates and correcting meiotic errors (Hong et al. 2018). In the future, it will be interesting to determine whether additional proteins assist in processing DNA tethers. In addition to its role in meiotic recombination, the BLM/HIM-6 helicase (which localizes to interfering CO sites, Woglar and Villeneuve 2018) has been shown in multiple systems to localize to and to help resolve ultrafine DNA bridges during both mitotic and meiotic chromosome segregation (Chan et al. 2007, Chan et al. 2009, Chan et al. 2018, Hong et al. 2016, McVey et al. 2007, Schvartzstein et al. 2014). Similarly, the nucleases MUS-81, SLX-1 and SLX-4 have genetic interactions with both LEM-3 (Hong and Velkova et al. 2018) and HIM-6 (Agostinho et al. 2013) and may act combinatorially to resolve DNA bridging due to excess COs. Further, topoisomerase II has been shown to remove DNA tethers from heterochromatin regions that connect achiasmate homologs in *Drosophila* (Hughes and Hawley 2014).

Although nucleases can serve as a generalized solution to resolve erroneous DNA connections, we also discovered a second mechanism specific to oocytes. In oocytes where a DNA tether was not resolved in anaphase I, the tethered DNA was preferentially extruded from the oocyte in anaphase II, presumably resulting in a euploid egg. This mechanism is made possible by the asymmetric nature of the oocyte divisions, in which half of the genetic material is discarded into the polar body during each division. This asymmetry has been shown to allow the preferential elimination of univalent chromosomes in *C. elegans* (Cortes et al. 2015), and here provides a second opportunity for error correction in the presence of excess COs. It is interesting to speculate on whether the oocyte spindle actively recognizes the DNA tether, or whether the tether passively biases the rotation of the spindle, positioning the tethered chromosome adjacent to the cortex, where it will be extruded into the second polar body. Currently, little is known about how spindle rotation is influenced in *C. elegans*; however, previous work has shown that the minus-end binding protein ASPM-1 is brighter on the spindle pole that rotates towards the cortex, suggesting that spindle asymmetry could play a role (Vargas et al. 2017) similar to spindle asymmetry and selfish centromeres in mouse oocytes (Akera et al. 2017). Going forward, it would be interesting to ask whether unresolved DNA connections influence spindle factors to allow for preferential spindle rotation as a form of meiotic drive.

Overall, our study provides new molecular insights into how excess COs affects chromosome congression and segregation in the oocyte. Further, our results indicate the existence of mechanisms to assist with correcting errors associated with the formation of excess COs, thereby lending important insight into how some organisms, such as *Saccharomyces cerevisiae*, can tolerate more than two interfering COs per chromosome. Future studies investigating why certain organisms can experience higher levels of interfering COs will lend further insight into these critical mechanisms for maintaining genomic integrity through generations.

## Materials and Methods

### C. elegans strains, genetics, and culture conditions

All strains are from the Bristol N2 background and were maintained at 15°C or 20°C and crossed at 20°C under standard conditions. Temperatures used for specific experiments are indicated below. For all experiments with meiotic mutants, homozygous mutant worms were derived from balanced heterozygous parents by selecting progeny lacking a dominant marker (Unc and/or GFP) associated with the balancer. For experiments marked 25°C, L4 worms were shifted to 25°C 24-48 hours prior to dissection.

The following strains were used in this study:

N2: Bristol wild-type strain.
AV311: *dpy-18(e364) unc-3(E151) meT7*(*III; X; IV*). (Hillers and Villeneuve 2003)
AV630: *meIs8[unc-119(+) Ppie-1::gfp::cosa-1] II*. (Yokoo et al. 2012)
OD868: *ItSi220*[pOD1249/pSW077; *Pmex-5::GFP-tbb-2-operon-linker-mCherry-his-11; cb-unc119(+)]I* (Wang et al. 2015)
DLW11: *meIs8[unc-119(+) Ppie-1::gfp::cosa-1] II; dpy-18(e364) unc-3(e151) meT7 (III;X;IV).* (This study)
DLW30: *ItSi220 I; dpy-18(e364) unc-3(E151) meT7(III; X; IV)* (This study)

### RNAi treatments

RNAi by feeding was performed as previously described (Muscat et al. 2015). Worms were synchronized at the L1 phase by bleaching adults and allowing resultant eggs to hatch on unseeded NGM plates at 20°C for 20-24 hrs. Synchronized L1s were then washed off of the unseeded NGM plates with M9 and placed on NGM+1mM IPTG+100μg ampicillin plates that were poured within 30 days of use and freshly seeded one day before use with clones picked from the Ahringer Lab RNAi feeding library (Kamath and Ahringer 2003) or the empty L4440 vector (referred to as “control RNAi” in figures and text). The RNAi plates with L1s were then placed at 15°C and grown to adulthood. For RNAi experiments performed at 25°C, L4 worms were shifted to 25°C for 40-48 hours prior to dissection.

### Immunofluorescence for late meiotic prophase I

Immunofluorescence was performed as in (Libuda et al. 2013). Gonads from adult worms at 18-24 hours post-L4 stage were dissected in 1x egg buffer with 0.1% Tween on VWR Superfrost Plus slides, fixed for 5 min in 1% paraformaldehyde, flash frozen with liquid nitrogen, and then fixed for 1 minute in 100% methanol at −20°C. Slides were washed 3 x 5 min in 1x PBST and blocked for one hour in 0.7% BSA in 1x PBST. Primary antibody dilutions were made in 1x PBST and added to slides. Slides were covered with a parafilm coverslip and incubated in a humid chamber overnight (14-18 hrs). Slides were washed 3 x 10 min in 1x PBST. Secondary antibody dilutions were made at 1:200 in 1x PBST using Invitrogen goat or donkey AlexaFluor labeled antibodies and added to slides. Slides were covered with a parafilm coverslip and placed in a humid chamber in the dark for 2 hrs. Slides were washed 3 x 10 min in 1x PBST in the dark. All washes and incubations were performed at room temperature, unless otherwise noted. 2 μg/ml DAPI was added to slides and slides were subsequently incubated in the dark with a parafilm coverslip in a humid chamber. Slides were washed once for 5 min in 1x PBST prior to mounting with Vectashield and a 20 x 40 mm coverslip with a 170 ± 5 μm thickness. Slides were sealed with nail polish immediately following mounting and then stored at 4°C prior to imaging. All slides were imaged (as described below) within two weeks of preparation. The following primary antibody dilutions were used: rabbit anti-GFP (1:1000) (Yokoo et al. 2012); chicken anti-GFP (1:1000) (Abcam 13970); guinea pig anti-SYP-1 (1:200) (MacQueen et al. 2002); rabbit anti-HIM-3 (1:200) (Zetka et al. 1999) and, chicken anti-HTP-3 (1:500) (MacQueen et al. 2005).

### Immunofluorescence for prometa-,meta- and anaphase I and II

Immunofluorescence was performed as in (Oogema et al. 2001). Briefly, adult worms were picked into a drop of M9 media on poly-l-lysine coated slides (Fisher scientific), cut in half to release oocytes, covered with a coverslip and frozen for 7 minutes in liquid nitrogen. The coverslip was quickly cracked off, and slides were fixed in −20°C methanol for 35 minutes. Samples were then rehydrated in PBS and blocked in AbDil (PBS with 4% BSA, 0.1% Triton X-100, 0.02% Sodium Azide) for one hour at room temperature, and then washed with PBST (PBS + 0.1% Triton X-100). Primary antibodies were diluted in AbDil, applied to samples and incubated overnight at 4°C. Samples were washed with PBST, and secondary antibodies were diluted in AbDil, applied to samples, and incubated for 2 hours at room temperature. Slides were washed and incubated with Hoechst 33342 (Invitrogen) at 1:1000 in PBST for 10 minutes at room temperature. Slides were washed a final time and mounted in 0.5% p-phenylenediamine in 90% glycerol, 20mM Tris, pH 8.8, sealed with nail polish, and stored at 4°C prior to imaging.

The following antibodies were used: rat anti-AIR-2 (1:500, Davis-Roca et al. 2017), rabbit anti-SEP-1 (1:200, gift from Andy Golden), rabbit anti-BIR-1 (1:800, this study), mouse anti-SUMO (1:500, gift from Federico Pelisch), rabbit anti-BUB-1 (1:1000, Davis-Roca et al. 2018), rabbit anti-ASPM-1 (1:5000, gift from Arshad Desai), Mouse anti-*α*-tubulin-FITC (1:500, DM1*α*, Sigma) and rabbit anti-KLP-19 (1:2500, this study). KLP-19 and BIR-1 polyclonal antibodies were generated by Covance using recombinant GST-BIR-1 (Full length protein) and GST-KLP-19 (amino acids 371-1084) as antigens (purification performed as in Davis-Roca et al. 2018). Antibody sera was then affinity purified and used at indicated concentrations. Alexa-fluor conjugated secondary antibodies were used at 1:500.

### Fixed Imaging

For Figure 1, IF slides were imaged at 512 x 512 pixel dimensions on an Applied Precision DeltaVision microscope using a 60x objective with 1.5x optivar. Images were acquired as Z-stacks at 0.2 μm intervals and deconvolved with Applied Precision softWoRx deconvolution software. For quantification of GFP::COSA-1 foci, nuclei that were in the last 4-5 rows of late pachytene and were completely contained within the image stack were analyzed. Foci were quantified manually from deconvolved three-dimensional stacks. *meT7* chromosomes in pachytene nuclei were identified based on size. Regardless of temperature (20°C or 25°C), all three wild-type/non-fused chromosomes in the nuclei of the *meT7* strains contained only one GFP::COSA-1 focus per chromosome (n=30), which is consistent with all chromosomes in wild-type strains and the wild-type/unfused chromosomes in *mnT12 (X;IV)* fusion chromosome strain (this study, Libuda et al. 2013, Yokoo et al. 2012). Given the consistency of COSA-1 counts for the 3 wild-type/non-fused chromosome in the *meT7* strains, the number of COSA-1 foci per *meT7* was calculated by subtracting 3 from the total number of COSA-1 foci per nucleus. For visualization and quantitation of chiasmata (Figures 1D), individual *meT7* bivalents from diakinesis nuclei in -2, -3, or -4 oocytes were identified based on size, cropped, and rotated in three-dimensions using Imaris (Bitplane/Oxford Instruments) three-dimensional rendering software. Scoring of chiasmata was based primarily on HTP-3 (chromosome axis) or HIM-3 (chromosome axis) and DAPI staining, as GFP::COSA-1 dissociates from chromosomes during progression through the diakinesis stage. For Figure 1B-C, images shown are projections through three-dimensional data stacks encompassing whole nuclei, generated with a maximum-intensity algorithm with the softWoRx (Applied Precision) software. For Figure 1D, the images of *meT7* bivalents shown are snapshots of a Imaris three-dimensional rendering of individual diakinesis bivalents with maximum intensity rendering for HTP-3. The images of wild type unfused autosome bivalents in Figure 1D are projections through three-dimensional data stacks encompassing the whole bivalent, generated with a maximum intensity algorithm with the softWoRx (Applied Precision) software.

For Figures 2, 3, 4, 5, and 7a, IF slides were imaged at 256 x 256 pixel dimensions on an Applied Precision DeltaVision microscope using a 100x oil objective (NA = 1.4), housed in the Northwestern Univeresity Biological Imaging Facility supported by the Northwestern University Office for Research. Images were acquired as Z-stacks at 0.2 μm intervals and deconvolved (ratio method, 15 cycles) with Applied Precision softWoRx deconvolution software. Images are maximum projections of entire spindles unless otherwise noted in the figure legends. Meiosis stages were determined by eye based on protein localization, chromosome-to-chromosome distance, chromosome size and polar body presence. *meT7* chromosomes were identified based on size.

### Ring Stretching Assay

Ring stretching on monopolar spindles in Figure 3 was performed as in (Muscat et al. 2015). Worms were grown on RNAi plates with a 1:1 mixture of *emb-30* and *klp-18* RNAi feeding clones from the Ahringer library to induce an arrest in metaphase on a monopolar spindle. RC components tend to stretch away from bivalents in extended metaphase arrest; spindles in extended arrest were determined by eye based on significant ring stretching of the three normal bivalents. The number of stretching rings on either *meT7* or a control normal bivalent was counted using Imaris.

### Quantification

#### Ring Structure Quantification

To assess ring structure, RCs on *meT7* bivalents were rotated in Imaris. Rings were scored by eye and considered mispatterned if they had more than one plane, consisted of more than one individual unit, or had more than one major loop. For Figure 2A, Rings were considered “slightly mispatterned” if they consisted of two distinct units on a single plane or appeared as two connected loops on a single plane. Rings with more than two units, multiple loops at different angles or planes on the bivalent, or rings with many fragments were considered “severely mispatterned.”

#### Linescans

For Figure 2, linescans were performed in ImageJ. Fluorescence intensity linescans of normal or fused chromosomes were performed along the pole-pole axis, as determined by tubulin intensity, at 40 x 30 pixels (L x W) (n = 25). Only clearly bipolar spindles were used for analysis, and all images had the same exposure conditions. Both chromosome length and fluorescence intensity were normalized to a maximum of 1, and the average (solid line) and standard error of the mean (SEM, shaded), of both DNA and SUMO were plotted using the ggplot2 package in Jupyter Notebook.

#### Metaphase Alignment Quantification

For Figure 4A, *meT7* worms were arrested in metaphase using *emb-30* RNAi. The metaphase plate was determined by rotating the image in 3D using Imaris until a single plane could be determined by eye based on SUMO-stained RCs on chromosomes I, II and V. The average total width of SUMO intensity of chromosomes I, II and V was measured using Imaris per spindle at the determined metaphase plate. The *meT7* fusion chromosome was considered “unaligned” if greater than 50% of its SUMO intensity fell outside of 2 standard deviations of the average width of chromosomes I, II and V from the determined metaphase plate.

#### Chromosome Distance Measurements

For Figure 4B, chromosome-to-pole distances were measured using Imaris. The center of the monopole was determined by using the “Surfaces” tool to determine the volume of the ASPM-1 region and to assign the center of this volume. Then, the distance between this point and the center of each chromosome was measured. Per spindle, the average of the distances to chromosomes I, II and V was subtracted from each individual distance to chromosomes I, II, or V (green points on the diagram/graph in Figure 3C) or to the fused chromosome (blue points). For Figure S3, “mid-to-late anaphase” was defined as having a chromosome-to-chromosome segregation distance of 2.5μM as measured by Imaris using the “surfaces” tool (Davis-Roca et al. 2017).

#### Chromatin Bridging Quantification

DNA was denoted as bridged if in anaphase the width of the bridge was less than the width at either end of the segregating chromosomes. Otherwise, a single *meT7* chromosome mass in anaphase was marked as showing delayed segregation, and two distinct segregating *meT7* masses with no connecting DNA were considered wild type.

#### Meiosis II Quatification

By eye, ploidy was determined based on chromosome counts on MII spindles (i.e. oocytes were considered aneuploid if the number of Hoechst-stained bodies was not equal to 4). Incomplete polar body extrusion was defined by a clear continual strand of DNA connecting a polar body to a chromatid pair within a forming or formed MII spindle. Polar bodies were determined to be DNA masses in close proximity to, yet remaining outside of, forming MII spindles.

### Scoring Embryonic Viability

Viability counts (percent hatching) were determined by singling out 5-15 L4s onto individual plates and growing them at 25°C until broods were produced. Mothers were transferred to new plates each day and allowed to produce broods until they no longer laid fertilized embryos. After mothers were moved, embryos and larvae were counted, and then returned to 25°C and allowed to develop for 18-24 hours (wild type) or 40-48 hours (*meT7*) before counting unhatched embryos. *meT7* embryos were given more time to develop, as the strain develops more slowly than wild type.

### Live Imaging

Live imaging of oocyte meiotic chromosome segregation in wild type and *meT7* fluorescent fusion lines was accomplished by cutting open adult worms with a single row or less of embryos in 4μl of egg buffer (118mM NaCl, 48mM KCl, 2mM CaCl_2_, 2mM MgCl_2_, and 0.025 mM HEPES, filter sterilized before HEPES addition) on a coverslip and gently mounting onto a 2% agarose pad on a microscope slide. Worms were synchronized by hypochlorite hatch-off, and plated worms were upshifted at 25°C overnight before imaging. Oocytes were imaged using a spinning disk confocal unit, CSU-W with Borealis (Andor), and dual iXon Ultra 897 (Andor) cameras mounted on an inverted Leica DMi8 microscope, with a 100x HCX PL APO 1.4NA oil objective lens (Leica). The imaging system was controlled via Metamorph (Molecular Devices) software. Oocytes were imaged every 5 seconds with 1μm Z-spacing (16μm total Z-stack) and the 488nm and 561nm channels were imaged simultaneously. After recording, movies were maximum projected, cropped, and color channels were adjusted independently for brightness and contrast in ImageJ (National Institutes of Health).

### Statistical analysis

All reported p-values are based on Pearson’s chi-square statistical tests comparing stated frequency distributions. A P-value less than 0.05 was considered statistically significant.

## Acknowledgements

We thank A. Davis-Roca and R. Ng for purifying the KLP-19 and BIR-1 antibodies, F. Pelisch, A. Golden, A. Dernburg and A. Villeneuve for antibodies and the CGC (funded by National Institutes of Health (NIH) P40 OD010440), A. Desai, and A. Villeneuve for strains. We thank K. Hillers and Wignall lab members N. Divekar, H. Horton, and I. Wolff for comments on the manuscript, and H. Horton for help with optimization of assays. This work was supported by the National Institutes of Health R01GM124354 to SMW, R00HD076165 and R35GM128890 to DEL, R01GM049869 and R35GM217221 to BB, T32GM007413 to AJS, and a Jane Coffin Childs Postdoctoral Fellowship to CKC. DEL is also a Searle Scholar and recipient of a March of Dimes Basil O’Connor Starter Scholar award.

## Figure Legends

**Supplemental Figure 1.**
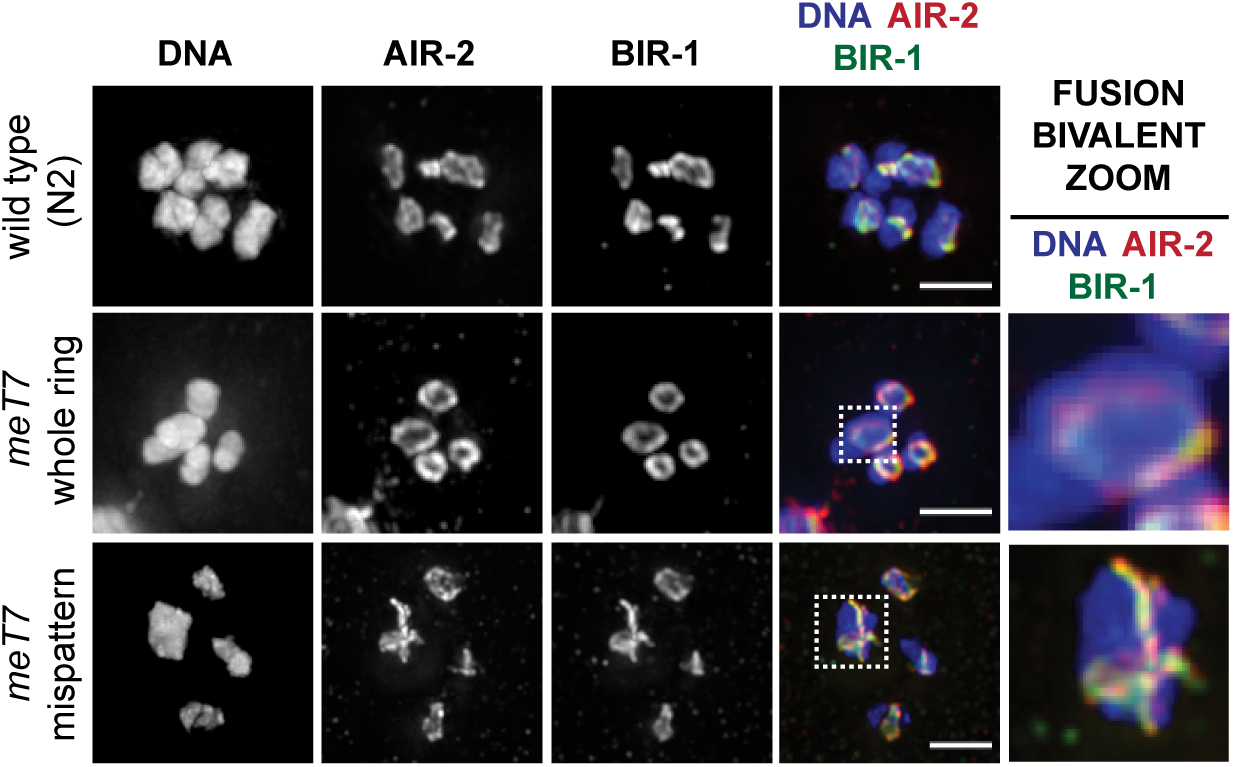
Additional characterization of CPC components (related to Figure 2). (A) BIR-1 (green) and AIR-2 (red) colocalize on all ring structure types on *meT7*. Scale bars=2.5µm.

**Supplemental Figure 2.**
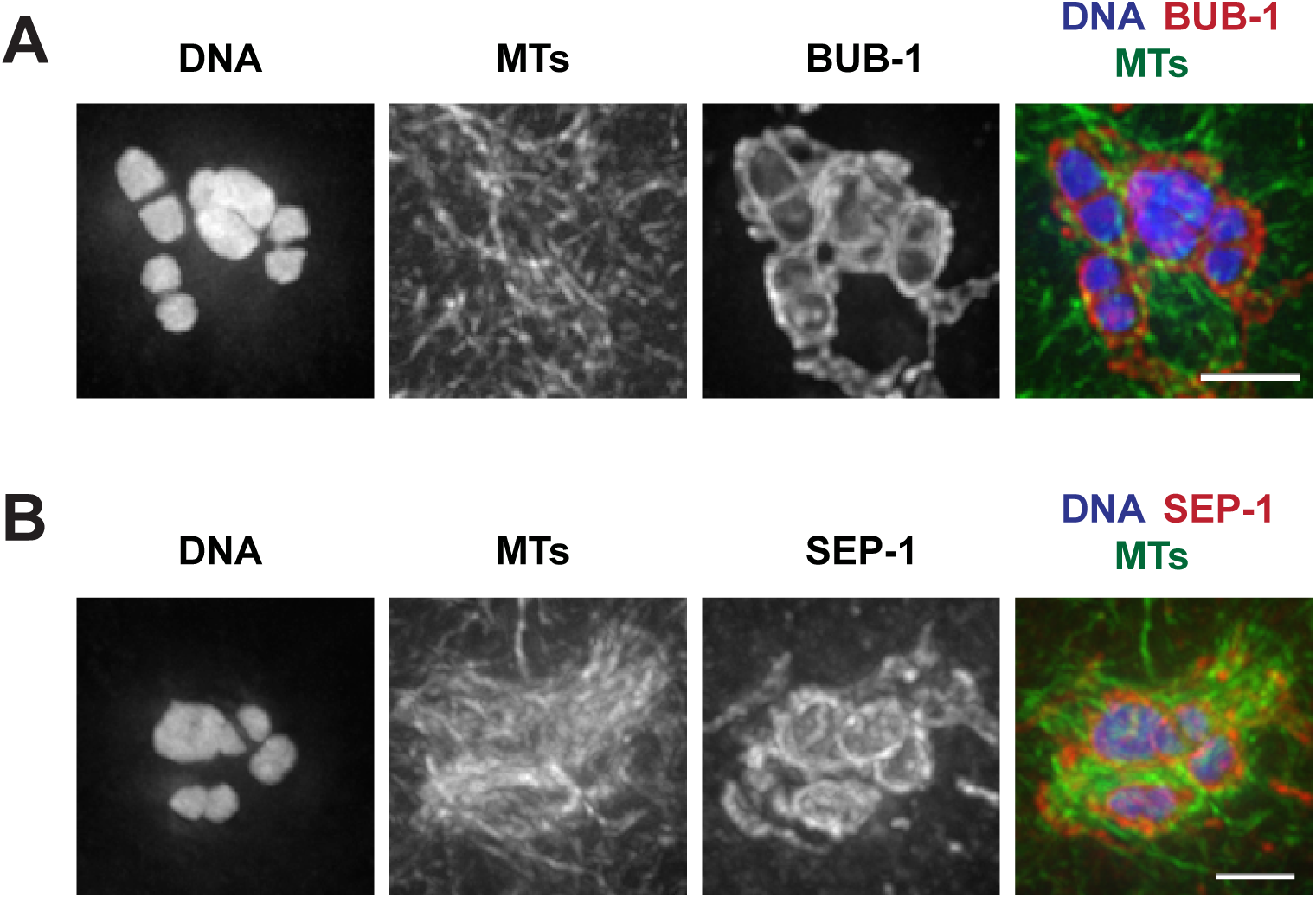
The holocentric kinetochore cups the entirety of *meT7*. (A) Localization of kinase BUB-1 (red) on prometaphase I chromosomes (blue); microtubules shown in green. Since *C. elegans* chromosomes are holocentric, kinetochore proteins cup the ends of normal bivalents, in addition to forming filaments within the spindle. In *meT7* fusion bivalents, the kinetochore cups the entirety of the bivalent. (B) Localization of SEP-1 (as another marker of the kinetochore, red) on prometaphase I chromosomes (blue), demonstrating the same organization as BUB-1. All scale bars=2.5µm.

**Supplemental Figure 3.**
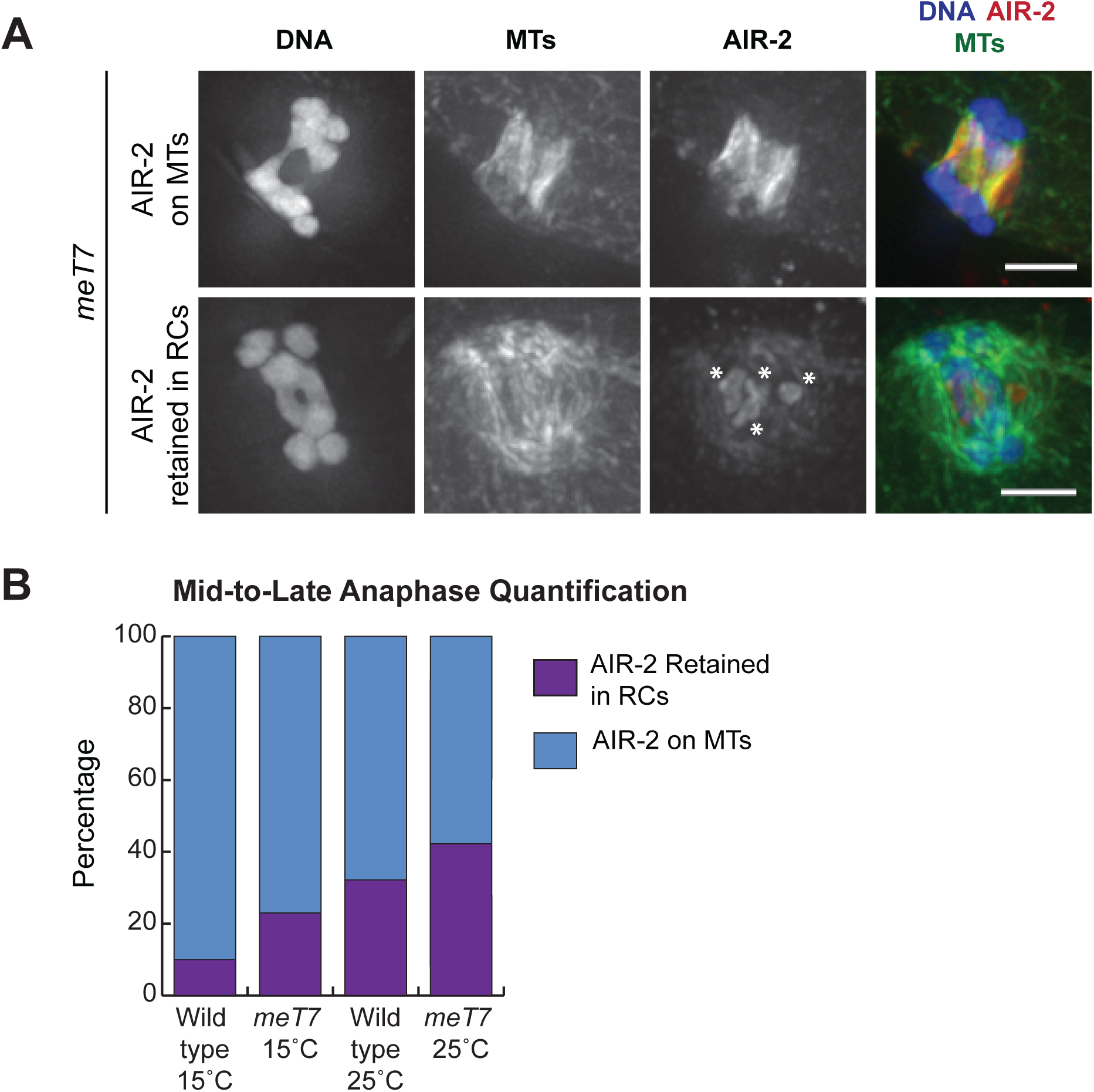
RCs show an increased frequency of delayed disassembly in *meT7* oocytes, demonstrating that errors are recognized. (A) In late anaphase, AIR-2 (red) either relocalizes to the spindle (top row) or stays RC-associated in *meT7* (bottom row; RCs indicated with asterisks), indicative of an RC disassembly delay. Scale bars=2.5µm. (B) Quantification of AIR-2 localization in late anaphase. At 15°C, 10% of N2 spindles showed a delay in ring disassembly, whereas 23% of *meT7* spindles showed the same behavior (N=31, both conditions). The frequency of RC delayed disassembly increased at 15°C to 33% in wild type (N=42) and 41% in meT7 spindles (N=29).

**Table S1.**
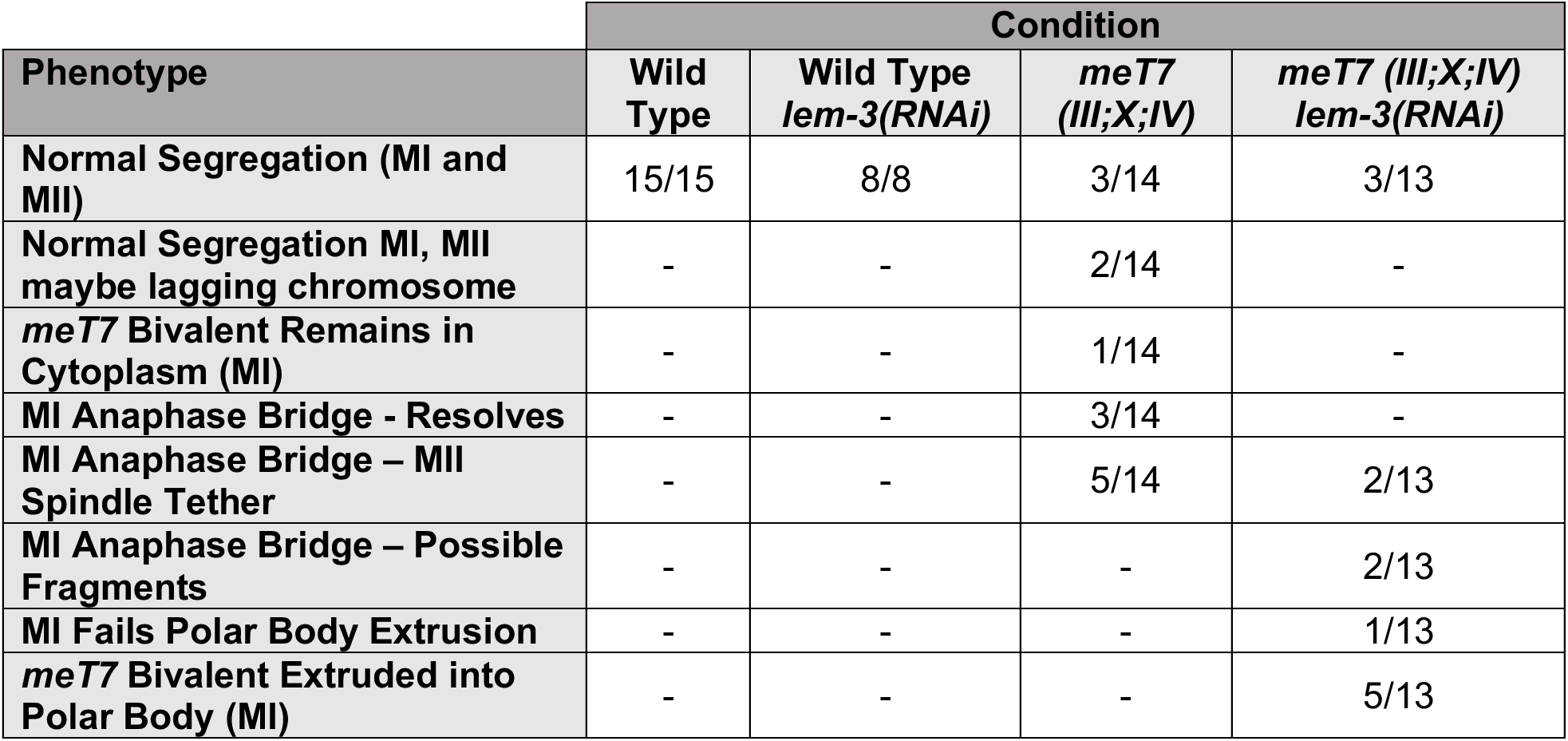
Frequency of chromosome segregation phenotypes from live imaging experiments.

**Supplemental Movie 1. Example movie showing *meT7* segregation occasionally (5 of 14 movies) results in the first polar body being tethered to the meiosis II spindle by a chromatin bridge (related to Figure 6 and Table S1).** Microtubules (GFP::TBB-2) shown in magenta and chromosomes (mCherry::HIS-11), shown in green, during both meiotic divisions. Playback framerate is 25 frames per second.

**Supplemental Movie 2. Example movie showing *meT7* failed to segregate in meiosis I and segregated with a chromatin bridge in meiosis II (1 of 14 movies) (related to Figure 6 and Table S1).** Microtubules (GFP::TBB-2) shown in magenta and chromosomes (mCherry::HIS-11) shown in green during both meiotic divisions. Playback framerate is 25 frames per second.

**Supplemental Movie 3 – Example movie showing *meT7* segregating with a chromatin bridge in meiosis I that appears to resolve before anaphase of meiosis II (3 of 14 movies) (related to Figure 6 and Table S1).** Microtubules (GFP::TBB-2) shown in magenta and chromosomes (mCherry::HIS-11) shown in green during both meiotic divisions. Playback framerate is 25 frames per second.

## References

Agostinho A, Meier B, Sonneville R, Jagut M, Woglar A, Blow J, Jantsch V, Gartner A. Combinatorial regulation of meiotic holliday junction resolution in C. elegans by HIM-6 (BLM) helicase, SLX-4, and the SLX-1, MUS-81 and XPF-1 nucleases. PLoS Genet. 2013;9(7):e1003591. doi: 10.1371/journal.pgen.1003591.

Akera T, Chmátal L, Trimm E, Yang K, Aonbangkhen C, Chenoweth DM, Janke C, Schultz RM, Lampson MA. Spindle asymmetry drives non-Mendelian chromosome segregation. Science. 2017 Nov 3;358(6363):668–672. doi: 10.1126/science.aan0092.

Chan KL, North PS, Hickson ID. BLM is required for faithful chromosome segregation and its localization defines a class of ultrafine anaphase bridges. EMBO J. 2007 Jul 25;26(14):3397–409. doi: 10.1038/sj.emboj.7601777.

Chan KL, Palmai-Pallag T, Ying S, Hickson ID. Replication stress induces sister-chromatid bridging at fragile site loci in mitosis. Nat Cell Biol. 2009 Jun;11(6):753–60. doi: 10.1038/ncb1882.

Chan YW, Fugger K, West SC. Unresolved recombination intermediates lead to ultra-fine anaphase bridges, chromosome breaks and aberrations. Nat Cell Biol. 2018 Jan;20(1):92–103. doi: 10.1038/s41556-017-0011-1.

Connolly AA, Sugioka K, Chuang CH, Lowry JB, Bowerman B. KLP-7 acts through the Ndc80 complex to limit pole number in C. elegans oocyte meiotic spindle assembly. J Cell Biol. 2015 Sep 14;210(6):917–32. doi: 10.1083/jcb.201412010.

Cortes DB, McNally KL, Mains PE, McNally FJ. The asymmetry of female meiosis reduces the frequency of inheritance of unpaired chromosomes. Elife. 2015 Apr 7;4:e06056. doi: 10.7554/eLife.06056.

Davis-Roca AC, Divekar NS, Ng RK, Wignall SM. Dynamic SUMO remodeling drives a series of critical events during the meiotic divisions in Caenorhabditis elegans. PLoS Genet. 2018 Sep;14(9):e1007626. doi: 10.1371/journal.pgen.1007626.

Davis-Roca AC, Muscat CC, Wignall SM. *Caenorhabditis elegans* oocytes detect meiotic errors in the absence of canonical end-on kinetochore attachments. J Cell Biol. 2017 May 1;216(5):1243–1253. doi: 10.1083/jcb.201608042.

Dumont J, Oegema K, Desai A. A kinetochore-independent mechanism drives anaphase chromosome separation during acentrosomal meiosis. Nat Cell Biol. 2010 Sep;12(9):894–901. doi: 10.1038/ncb2093.

Gray S, Cohen PE. Control of Meiotic Crossovers: From Double-Strand Break Formation to Designation. Annu Rev Genet. 2016 Nov 23;50:175–210. doi: 10.1146/annurev-genet-120215-035111.

Han X, Adames K, Sykes EM, Srayko M. The KLP-7 Residue S546 Is a Putative Aurora Kinase Site Required for Microtubule Regulation at the Centrosome in C. elegans. PLoS One. 2015;10(7):e0132593. doi: 10.1371/journal.pone.0132593.

Hillers KJ, Villeneuve AM. Chromosome-wide control of meiotic crossing over in C. elegans. Curr Biol. 2003 Sep 16;13(18):1641–7.

Hong Y, Sonneville R, Wang B, Scheidt V, Meier B, Woglar A, Demetriou S, Labib K, Jantsch V, Gartner A. LEM-3 is a midbody-tethered DNA nuclease that resolves chromatin bridges during late mitosis. Nat Commun. 2018 Feb 20;9(1):728. doi: 10.1038/s41467-018-03135-w.

Hong Y, Velkova M, Silva N, Jagut M, Scheidt V, Labib K, Jantsch V, Gartner A. The conserved LEM-3/Ankle1 nuclease is involved in the combinatorial regulation of meiotic recombination repair and chromosome segregation in Caenorhabditis elegans. PLoS Genet. 2018 Jun;14(6):e1007453. doi: 10.1371/journal.pgen.1007453.

Hong Y, Sonneville R, Agostinho A, Meier B, Wang B, Blow JJ, Gartner A. The SMC-5/6 Complex and the HIM-6 (BLM) Helicase Synergistically Promote Meiotic Recombination Intermediate Processing and Chromosome Maturation during Caenorhabditis elegans Meiosis. PLoS Genet. 2016 Mar;12(3):e1005872. doi: 10.1371/journal.pgen.1005872.

Howe M, McDonald KL, Albertson DG, Meyer BJ. HIM-10 is required for kinetochore structure and function on Caenorhabditis elegans holocentric chromosomes. J Cell Biol. 2001 Jun 11;153(6):1227–38. doi: 10.1083/jcb.153.6.1227.

Hughes SE, Hawley RS. Topoisomerase II is required for the proper separation of heterochromatic regions during Drosophila melanogaster female meiosis. PLoS Genet. 2014 Oct;10(10):e1004650. doi: 10.1371/journal.pgen.1004650.

Kaitna S, Pasierbek P, Jantsch M, Loidl J, Glotzer M. The aurora B kinase AIR-2 regulates kinetochores during mitosis and is required for separation of homologous chromosomes during meiosis. Curr Biol. 2002 May 14;12(10):798–812. doi: 10.1016/s0960-9822(02)00820-5.

Kamath RS, Ahringer J. Genome-wide RNAi screening in Caenorhabditis elegans. Methods. 2003 Aug;30(4):313–21. doi: 10.1016/s1046-2023(03)00050-1.

Laband K, Le Borgne R, Edwards F, Stefanutti M, Canman JC, Verbavatz JM, Dumont J. Chromosome segregation occurs by microtubule pushing in oocytes. Nat Commun. 2017 Nov 14;8(1):1499. doi: 10.1038/s41467-017-01539-8.

Libuda DE, Uzawa S, Meyer BJ, Villeneuve AM. Meiotic chromosome structures constrain and respond to designation of crossover sites. Nature. 2013 Oct 31;502(7473):703–6. doi: 10.1038/nature12577.

MacQueen AJ, Colaiácovo MP, McDonald K, Villeneuve AM. Synapsis-dependent and - independent mechanisms stabilize homolog pairing during meiotic prophase in C. elegans. Genes Dev. 2002 Sep 15;16(18):2428–42. doi: 10.1101/gad.1011602.

MacQueen AJ, Phillips CM, Bhalla N, Weiser P, Villeneuve AM, Dernburg AF. Chromosome sites play dual roles to establish homologous synapsis during meiosis in C. elegans. Cell. 2005 Dec 16;123(6):1037–50. doi: 10.1016/j.cell.2005.09.034.

Martinez-Perez E, Colaiácovo MP. Distribution of meiotic recombination events: talking to your neighbors. Curr Opin Genet Dev. 2009 Apr;19(2):105–12. doi: 10.1016/j.gde.2009.02.005.

Martinez-Perez E, Schvarzstein M, Barroso C, Lightfoot J, Dernburg AF, Villeneuve AM. Crossovers trigger a remodeling of meiotic chromosome axis composition that is linked to two-step loss of sister chromatid cohesion. Genes Dev. 2008 Oct 15;22(20):2886–901. doi: 10.1101/gad.1694108.

McVey M, Andersen SL, Broze Y, Sekelsky J. Multiple functions of Drosophila BLM helicase in maintenance of genome stability. Genetics. 2007 Aug;176(4):1979–92. doi: 10.1534/genetics.106.070052.

Melters DP, Paliulis LV, Korf IF, Chan SW. Holocentric chromosomes: convergent evolution, meiotic adaptations, and genomic analysis. Chromosome Res. 2012 Jul;20(5):579–93. doi: 10.1007/s10577-012-9292-1.

Monen J, Maddox PS, Hyndman F, Oegema K, Desai A. Differential role of CENP-A in the segregation of holocentric C. elegans chromosomes during meiosis and mitosis. Nat Cell Biol. 2005 Dec;7(12):1248–55. doi: 10.1038/ncb1331.

Muscat CC, Torre-Santiago KM, Tran MV, Powers JA, Wignall SM. Kinetochore-independent chromosome segregation driven by lateral microtubule bundles. Elife. 2015 May 30;4:e06462. doi: 10.7554/eLife.06462.

Mullen TJ, Wignall SM. Interplay between microtubule bundling and sorting factors ensures acentriolar spindle stability during C. elegans oocyte meiosis. PLoS Genet. 2017 Sep; 13(9):e1006986. doi: 10.1371/journal.pgen.1006986.

Muller H. The Mechanism of Crossing-Over. The American Naturalist. 1916. 50(592), 193–221.

Nabeshima K, Villeneuve AM, Colaiácovo MP. Crossing over is coupled to late meiotic prophase bivalent differentiation through asymmetric disassembly of the SC. J Cell Biol. 2005 Feb 28;168(5):683–9. doi: 10.1083/jcb.200410144.

Nokkala S, Kuznetsova VG, Maryanska-Nadachowska A, Nokkala C. Holocentric chromosomes in meiosis. I. Restriction of the number of chiasmata in bivalents. Chromosome Res. 2004;12(7):733–9. doi: 10.1023/B:CHRO.0000045797.74375.70.

Oegema K, Desai A, Rybina S, Kirkham M, Hyman AA. Functional analysis of kinetochore assembly in Caenorhabditis elegans. J Cell Biol. 2001 Jun 11;153(6):1209–26. doi: 10.1083/jcb.153.6.1209.

Pelisch F, Bel Borja L, Jaffray EG, Hay RT. Sumoylation regulates protein dynamics during meiotic chromosome segregation in C. elegans oocytes. J Cell Sci. 2019 Jul 18;132(14). doi: 10.1242/jcs.232330.

Pelisch F, Tammsalu T, Wang B, Jaffray EG, Gartner A, Hay RT. A SUMO-Dependent Protein Network Regulates Chromosome Congression during Oocyte Meiosis. Mol Cell. 2017 Jan 5;65(1):66–77. doi: 10.1016/j.molcel.2016.11.001.

Rogers E, Bishop JD, Waddle JA, Schumacher JM, Lin R. The aurora kinase AIR-2 functions in the release of chromosome cohesion in Caenorhabditis elegans meiosis. J Cell Biol. 2002 Apr 15;157(2):219–29. doi: 10.1083/jcb.200110045.

Romano A, Guse A, Krascenicova I, Schnabel H, Schnabel R, Glotzer M. CSC-1: a subunit of the Aurora B kinase complex that binds to the survivin-like protein BIR-1 and the incenp-like protein ICP-1. J Cell Biol. 2003 Apr 28;161(2):229–36. doi: 10.1083/jcb.200207117.

Schumacher JM, Golden A, Donovan PJ. AIR-2: An Aurora/Ipl1-related protein kinase associated with chromosomes and midbody microtubules is required for polar body extrusion and cytokinesis in Caenorhabditis elegans embryos. J Cell Biol. 1998 Dec 14;143(6):1635–46. doi: 10.1083/jcb.143.6.1635.

Schvarzstein M, Pattabiraman D, Libuda DE, Ramadugu A, Tam A, Martinez-Perez E, Roelens B, Zawadzki KA, Yokoo R, Rosu S, Severson AF, Meyer BJ, Nabeshima K, Villeneuve AM. DNA helicase HIM-6/BLM both promotes MutSγ-dependent crossovers and antagonizes MutSγ-independent interhomolog associations during caenorhabditis elegans meiosis. Genetics. 2014 Sep;198(1):193–207. doi: 10.1534/genetics.114.161513.

Siomos MF, Badrinath A, Pasierbek P, Livingstone D, White J, Glotzer M, Nasmyth K. Separase is required for chromosome segregation during meiosis I in Caenorhabditis elegans. Curr Biol. 2001 Nov 27;11(23):1825–35. doi: 10.1016/s0960-9822(01)00588-7.

Stimpson KM, Matheny JE, Sullivan BA. Dicentric chromosomes: unique models to study centromere function and inactivation. Chromosome Res. 2012 Jul;20(5):595–605. doi: 10.1007/s10577-012-9302-3. Review.

Sturtevant AH. The linear arrangement of six sex-linked factors in Drosophila, as shown by their mode of association. J. Exp. Zool. 1913. 14: 43–59. doi:10.1002/jez.1400140104

Vargas E, McNally K, Friedman JA, Cortes DB, Wang DY, Korf IF, McNally FJ. Autosomal Trisomy and Triploidy Are Corrected During Female Meiosis in *Caenorhabditis elegans*. Genetics. 2017 Nov;207(3):911–922. doi: 10.1534/genetics.117.300259.

Wang S, Hassold T, Hunt P, White MA, Zickler D, Kleckner N, Zhang L. Inefficient Crossover Maturation Underlies Elevated Aneuploidy in Human Female Meiosis. Cell. 2017 Mar 9;168(6):977–989.e17. doi: 10.1016/j.cell.2017.02.002.

Wang S, Wu D, Quintin S, Green RA, Cheerambathur DK, Ochoa SD, Desai A, Oegema K. NOCA-1 functions with γ-tubulin and in parallel to Patronin to assemble non-centrosomal microtubule arrays in C. elegans. Elife. 2015 Sep 15;4:e08649. doi: 10.7554/eLife.08649.

Wendel JF, Greilhuber J, Leitch I, Dolezel J. (2012). Plant genome diversity. 10.1007/978-3-7091-1130-7.

Wignall SM, Villeneuve AM. Lateral microtubule bundles promote chromosome alignment during acentrosomal oocyte meiosis. Nat Cell Biol. 2009 Jul;11(7):839–44. doi: 10.1038/ncb1891.

Woglar A, Villeneuve AM. Dynamic Architecture of DNA Repair Complexes and the Synaptonemal Complex at Sites of Meiotic Recombination. Cell. 2018 Jun 14;173(7):1678–1691.e16. doi: 10.1016/j.cell.2018.03.066.

Wolff ID, Tran MV, Mullen TJ, Villeneuve AM, Wignall SM. Assembly of Caenorhabditis elegans acentrosomal spindles occurs without evident microtubule-organizing centers and requires microtubule sorting by KLP-18/kinesin-12 and MESP-1. Mol Biol Cell. 2016 Oct 15;27(20):3122–31. doi: 10.1091/mbc.E16-05-0291.

Yokoo R, Zawadzki KA, Nabeshima K, Drake M, Arur S, Villeneuve AM. COSA-1 reveals robust homeostasis and separable licensing and reinforcement steps governing meiotic crossovers. Cell. 2012 Mar 30;149(1):75–87. doi: 10.1016/j.cell.2012.01.052.

Zetka MC, Kawasaki I, Strome S, Müller F. Synapsis and chiasma formation in Caenorhabditis elegans require HIM-3, a meiotic chromosome core component that functions in chromosome segregation. Genes Dev. 1999 Sep 1;13(17):2258–70. doi: 10.1101/gad.13.17.2258.

